# Evidence for balancing selection on loci associated with host use in the genome of a generalist plant parasite

**DOI:** 10.1101/2025.04.02.646902

**Authors:** McCall B. Calvert, Alison Chen, Adya Gupta, Linda Wu, Corlett W. Wood

**Affiliations:** Department of Biology, University of Pennsylvania, Philadelphia, PA, USA

**Author notes:** Correspondence: M.C., C.W.W.

**Keywords:** Generalist, host-parasite interaction, heterogeneous selection, *Meloidogyne hapla*, root knot nematode, balancing selection

## Abstract

Most research on host-parasite interactions has focused on tightly coevolved species pairs, yet many parasites are generalists that infect a wide range of host taxa. Generalist parasites are subject to highly variable selection pressures across different hosts and the footprints that this heterogeneous selection leaves in the genomes of generalists have been largely unexplored. To address this knowledge gap, we studied patterns of host-associated genetic diversity in the generalist plant parasite, the northern root knot nematode *Meloidogyne hapla*. By leveraging whole genome amplification, we generated whole genome libraries of individual root-knot nematodes collected from three different host plant species in a spatially replicated design. Despite finding no evidence of host-associated population structure, we did observe several hundred host-associated SNPs putatively under differential host selection. Genomic regions containing these host-associated SNPs were enriched for signals of balancing selection. Additionally, host-associated SNPs were enriched for functions involved in plant pathogenesis, specifically the modification of plant cell walls. Host-associated SNPs were also found in genes with known roles in dampening host plant immune responses to infection. Together, our results suggest a model of balancing selection in which heterogeneous selection by different host taxa maintains adaptive diversity at loci involved in pathogenesis.

## Introduction

Hosts impose strong selection on parasites. This strong selection maintains high amounts of diversity in pathogen populations, particularly in genetic loci involved in virulence (Ebert and Fields 2020). The bulk of theoretical and empirical work on host-parasite interactions has focused on tightly coevolved pairs (Clarke 1976; Jaenike 1978; Hamilton 1980; Ebert 2008). This work has led to well-developed theory and clear predictions on the evolutionary dynamics of coevolved interactions (Lively 1987; Dybdahl and Lively 1998; Paterson et al. 2010; Morran et al. 2011; Thrall et al. 2012; Papkou et al. 2019). Yet, many host-parasite interactions lack this tight specificity: hosts are attacked by multiple parasite species and most parasites can invade multiple host species (Woolhouse et al. 2001; Power and Mitchell 2004; Barrett et al. 2009; Gibson 2019).

The majority of parasite taxa are generalists (Barrett et al. 2009). The heterogeneous selection pressure exerted by different host species can either favor the evolution of broadly-adapted, generalist genotypes or multiple specialist genotypes (Levins 1968; Felsenstein 1976; Kassen 2002; Bono et al. 2017; Visher and Boots 2020). To date, population genetic studies of generalists have been primarily focused on whether these species are true generalists or are composed of distinct host-associated populations (i.e. host ecotypes) (Cole and Viney 2018; Montarry et al. 2021). Such studies have historically relied on small marker sets, such as microsatellites or mitochondrial DNA, to do so.

However, even in the genomes of true generalists, some regions may be subject to conflicting host selection pressures. Whenever different alleles are favored on different hosts, such regions should carry signals of balancing selection. The results of experimental evolution studies in the lab that impose heterogeneous selection are consistent with this prediction (Kassen 2002; Bono et al. 2017). However, the extent of host-associated genetic diversity in wild populations of generalist parasites is still underexplored (but see (Vidal et al. 2019)). This is largely because parasites are small, challenging to isolate, and difficult to rear in the lab, making individual-level whole genome data from natural populations hard to collect. Furthermore, the small marker sets that have historically been applied to wild generalist parasites are insufficient to detect genomic signatures of selection. Characterizing the genomic signatures of differential selection across host species in a generalist parasite is crucial to understand the evolutionary dynamics of diffuse host-parasite interactions.

In this study, we leveraged whole genome amplification of individual microparasites to interrogate the genomic signatures of selection in a generalist parasite of plants: the northern root-knot nematode (*Meloidogyne hapla*). *M*. *hapla* is a destructive agricultural pest and is one of the most extreme generalists known to science, with over 80 documented host plant families (Jones et al. 2013; Ferris et al. 2024). *M. hapla* invades host plants via root tips and migrates to a desirable feeding site near the primary phloem (Eisenback and Triantaphyllou 1991). Once a feeding site is found, *M. hapla* penetrate host cells with their stylets and inject effectors that suppress plant immune responses, act as plant hormone mimics, and modify plant cell walls (Castagnone-Sereno et al. 2013; Mitchum and Liu 2022). These effectors also facilitate the de-differentiation of host cells into enlarged multinucleate “giant cells” which supply nematodes with their nutrient requirements and lead to the formation of visible root galls (Eisenback and

Triantaphyllou 1991; Mitchum and Liu 2022; Bali and Gleason 2024). Genotypes of *M. hapla* exhibit little variation in the breadth of host species they are capable of infecting, but do exhibit variation in performance across host species (Liu and Williamson 2006). Like other root knot nematode species, *M. hapla* possesses limited dispersal ability and therefore has little control over the host species they encounter (Prot 1980). As a result, it is likely that *M. hapla* experiences heterogeneous selection by different host species across space and through time.

In this study, we re-sequenced the genomes of 78 *M. hapla* individuals collected from root galls on field-sampled plants in a spatially replicated design. Specifically, we sampled *M. hapla* from three different host species, *Medicago lupulina*, *Trifolium repens*, and *Leucanthemum vulgare*, at each of four sampling sites (Fig. 1 A). By comparing the genomes of individuals collected from different host plants, we addressed the following questions: (1) Is there evidence for host-specialized ecotypes? (2) Is there evidence for adaptive genetic variation involved in host adaptation? (3) Are loci putatively involved in differential host adaptation predominantly under positive or balancing selection? And (4) Are loci putatively involved in differential host adaptation enriched for molecular functions involved in plant infection? We also explored patterns of direct selection on *Meloidogyne* genes previously demonstrated to be involved in plant infection. We found no evidence for host-associated ecotypes. However, *M. hapla* populations still contain extensive host-associated genetic variation. Host-associated loci are enriched for signals of balancing selection and regions under balancing selection tended to be older than the rest of the genome. Finally, host-associated loci are enriched for molecular functions likely involved in the modification of plant cell walls, which is a key process in the formation of their feeding sites within galls. Together, our results suggest that balancing selection driven by heterogeneous selection by different host species maintains variation in the genome of this generalist parasite.

**Figure 1:**
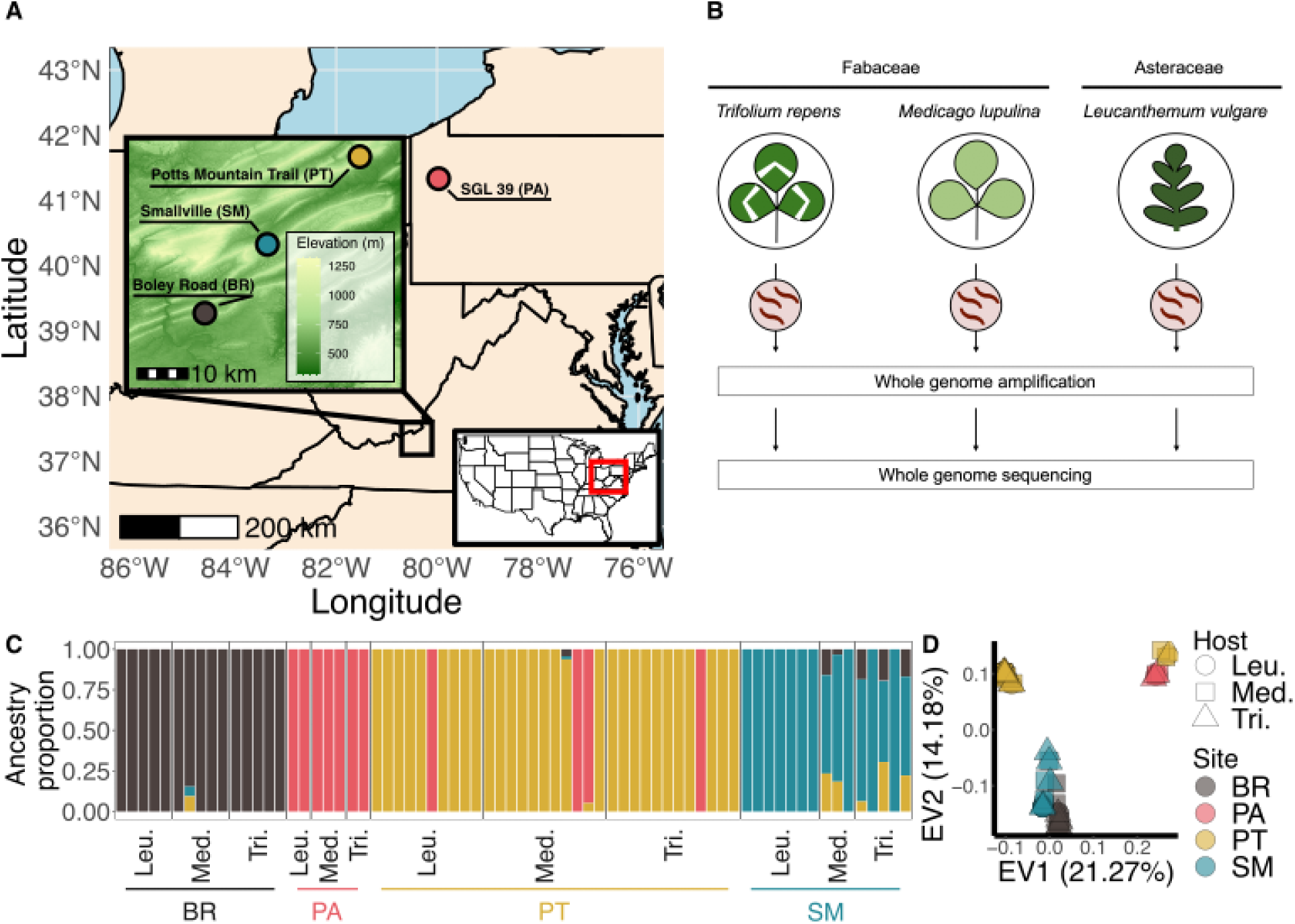
Sampling locations, sampling design, and population structure for the root knot nematodes collected in this study. (A) Sampling locations in Virgina and Pennsylvania. (B) Sampling design for this study. Root-knot nematode infected *Tritium repens, Medicago lupulina,* and *Leucanthemum vulgare* individuals were collected at each sampling location. Whole genome amplification followed by whole genome sequencing was performed on individual root-knot nematodes dissected from host plant roots. (C) Ancestry proportions for each individual root-knot nematode calculated using fastSTRUCTURE with *K = 4* and colored by the dominant cluster at each sampling location. Each bar represents an individual root knot nematode and are grouped by the host plant species they were sampled from and sampling location. (D) PCA showing the first and second eigenvectors of genetic variation in the root-knot nematodes sampled. Points represent individuals and are colared by sampling location according to the dominant population clusters identified in the fastSTRUCTURE analysis. Point shapes correspond the host plant species that individuals were sample from. In panels (C) and (D), host plant species are abbreviated; Leu. = *Leucanthemum vulgare,* Med. = *Medicago lupulina,* and Tri. = *Trifolium repens*.

## Results

Whole genome libraries for individual nematodes were produced using whole genome amplification (Fig. 1B). After alignment to the existing *M. hapla* reference assembly (Opperman et al. 2008), 13 samples that had fewer than 80% of reads mapping to the genome were removed from the dataset. After variant calling, and standard variant filtering (see Materials and Methods), we produced a set of 117,218 biallelic SNPs for downstream analysis.

### *Geography, but not host species, drives population structure in* M. hapla

We observed a strong effect of geography, but not host species, on *M. hapla* population structure (Fig. 1C,D). A fastSTRUCTURE (Raj et al. 2014) analysis found that the most likely number of population clusters was *K*=4, matching the number of sampling locations. All individuals within each sampling location showed complete or majority assignment to the dominant population cluster in that location except for four samples from PT, which were assigned to the same dominant population cluster as individuals from PA (Fig. 1C). Phylogenetic analysis of de-novo assembled mitochondrial genomes suggests that the PA samples and the four aforementioned PT individuals may belong to a different cytological type of *M. hapla* (Supplementary Figure S1). A PCA corroborated the findings from fastSTRUCURE (Fig. 1D). To explore whether the large genetic differences between the PA population and the three VA populations could be concealing a signal of host-associated population structure in Virginia, we also performed fastSTRUCTURE and PCA on the 59 individuals collected from sites in Virginia (excluding the aforementioned four individuals from PT). We again observed a strong effect of geography and no effect of host species (Supplementary Figure S2).

### Genome-wide scans reveal host-associated SNPs

We employed two complementary analyses to identify host-associated SNPs, both using an outlier approach implemented in the genome-environment association analysis tool, BayPass (Gautier 2015; Olazcuaga et al. 2020). In our first analysis, we leveraged all 59 samples across the three sampling sites in Virginia to construct a list of globally host-associated SNPs while controlling for population structure. In our second approach, we used only the 29 samples collected at the most well-sampled site, PT, to construct a list of host-associated SNPs local to that site. Each of these approaches involved making three comparisons between each pair of nematode samples collected from each host species: nematodes collected from *L. vulgare* and *M. lupulina* (Leu. vs. Med.), nematodes collected from *L. vulgare* and *T. repens* (Leu. vs. Tri.), and nematodes collected from *T. repens* and *M. lupulina* (Tri. vs. Med.).

When leveraging samples from all three sampling sites, we found a global set of 2,010 host-associated SNPs, with absolute allele frequency differences ranging between 0.08 and 0.21, depending on the host-associated nematode sample comparison (Fig. 2A,C,E). The results were qualitatively similar when we averaged the allele frequency differences of host-associated SNPs in non-overlapping 10kb windows, to control for physical linkage (Fig. 2A,C).

**Figure 2:**
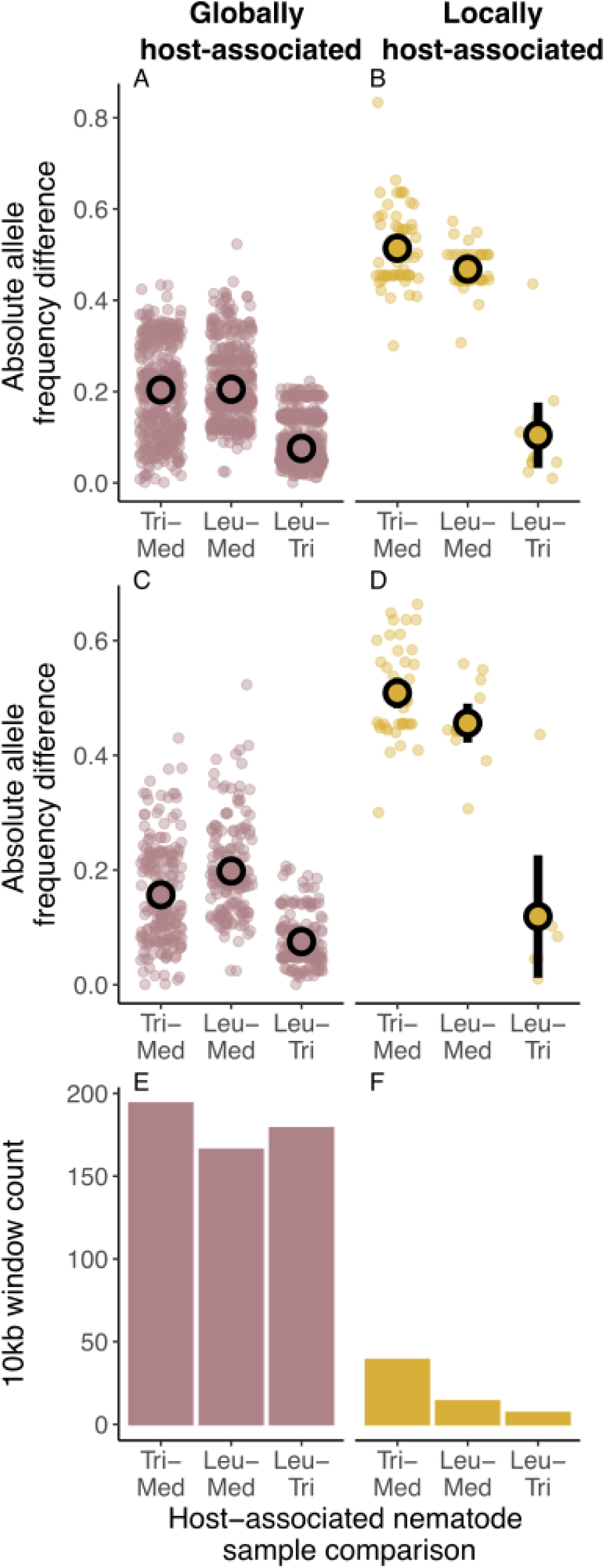
Absolute allele frequency differences of significant host-associated SNPs identified with BayPass using samples from (A,C,E) all sites (globally host-associated) and (B,D,F) using samples from only the PT site (locally host-associated). Panels A and B display the observed absolute allele frequency differences for globally host-associated and locally host-associated SNPs, respectively. Panels C and D plot the absolute allele frequency differences averaged by 10KB non-overlapping windows for globally host-associated and locally host-associated SNPs, respectively. Panels E and F display the total count of 1Okb windows harboring globally host-associated and locally host-associated SNPs, respectively.

Employing BayPass locally at PT revealed a total of 87 locally host-associated SNPs (Fig. 2B,D,F). Of these 87 SNPs, 12 were also present in the list of globally host-associated SNPs, which was significantly more than expected by chance (OR=10.63, P<0.0001). Furthermore, population structure-corrected standardized allele frequency differences were strongly correlated between the global and local analyses (Supplementary Figure S3). The average allele frequency difference for host-associated SNPs identified locally at PT tended to be greater than globally host-associated SNPs, ranging from 0.10 to 0.51, and, when averaging across 10kb windows, from 0.12 to 0.51 (Fig. 2B,D).

### Host-associated SNPs are enriched for balancing selection

To determine whether any host-associated SNPs are putatively under positive or balancing selection, we quantified Tajima’s *D* in 10 kb, non-overlapping windows in our most well-sampled site, PT (see Methods). Positive values of Tajima’s *D* indicate an excess of intermediate-frequency polymorphisms, consistent with balancing selection; negative values of Tajima’s *D* indicate an excess of low frequency polymorphisms, consistent with a recent selective sweep (i.e. positive selection). Positive and negative Tajima’s *D* values can also be indicative of recent population expansions or contractions, respectively (Tajima 1989). Our demographic inference using coalescent simulations in fastsimcoal found very little population growth (Supplementary Table S1), suggesting population size changes have a minimal effect on observed Tajima’s *D* values.

We found that 10kb windows containing host-associated SNPs had higher Tajima’s D values that the rest of the genome, slightly so for 10kb windows with globally host-associated SNPs (*F_1,2905_* = 1.812, P=0.178), and significantly so for 10kb windows with host-associated SNPs identified locally at PT (via permutation tests; *F_1,2905_* = 24.631 P<0.0001) (Fig 3 A,C).

**Figure 3:**
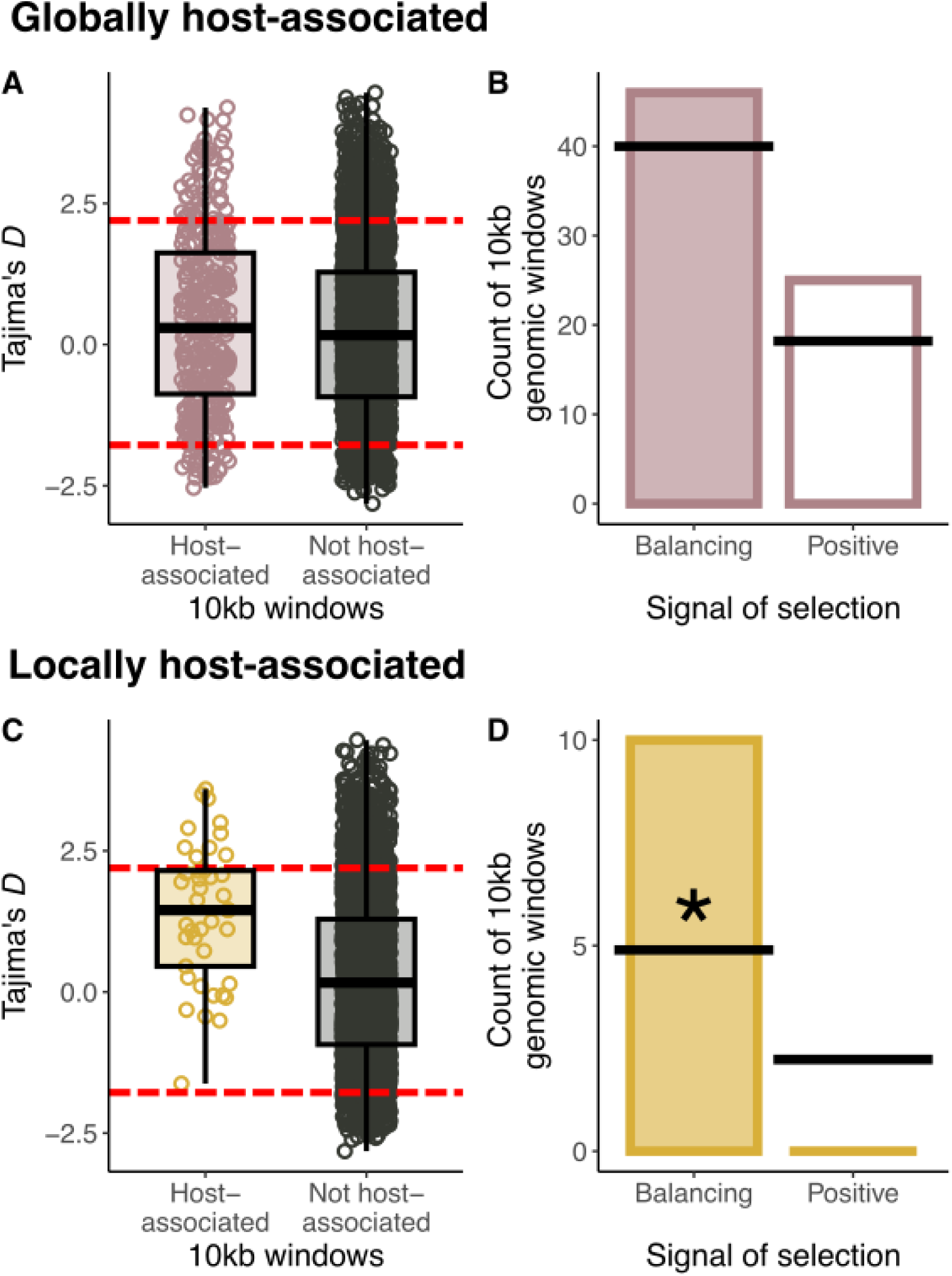
Signals of selection in genomic windows harboring (A.B) globally host-associated SNPs and (C.D) locally host-associated SNPs compared to the rest of the genome. Distributions of Tajima’s *D* values are plotted for 10 kb windows that do versus do not contain (A) globally host-associated SNPs and (C) locally host-associated SNPs. In these panels, each point represents the Tajima’s *D* value for a single 10 kb window. Dashed red lines represented the empirical 0.5% and 99.5% significance thresholds for determining the significance of a signal for selection using simulated genetic data. Tests for enrichment of significantly positive (balancing selection) and significantly negative (positive selection) Tajima’s *D* values in genomic windows that contain globally host-associated SNPs and locally host-associated SNPs are shown in panels B and D, respectively. Panels B and D contain bar plots of the observed number of genomic windows that contain host associated SNPs and exhibit a significant signal of balancing (shaded bar) or positive selection (open bar). Black lines mark the expected number of genomic windows in a given category and stars indicate significant enrichment tested using a fisher exact test.

We then tested for enrichment of host-associated SNPs in regions that exhibited signals of selection. To threshold for significant signals of selection, we used the 0.5% and 95.5% quantiles of a distribution of Tajima’s *D* values calculated from genetic data simulated from a best-fitting demographic model (see Methods). 10kb windows exhibiting significantly positive values of Tajima’s D were more common than those with significantly negative values. However, only 10kb windows containing host-associated SNPs identified locally at PT were significantly enriched for significant positive values of Tajima’s *D* (OR = 2.41, P = 0.02; Fig 3, B,D).

### Genomic windows with signals of balancing selection are older than putatively neutral regions

Balancing selection is expected to maintain selected alleles at intermediate frequencies for extended periods of time. Therefore, regions under balancing selection should be older than regions under positive selection. We used ARGweaver (Rasmussen et al. 2014) to estimate the time to the most recent common ancestor (TMRCA) for SNPs averaged across the same 10kb windows used in the Tajima’s *D* analysis. To avoid confounding effects of population structure, TMRCAs were estimated using only samples from our most well-sampled site, PT. Owing to the fragmented status of the *M. hapla* reference assembly, our inference on TMRCA was limited to contigs for which the ARGweaver models successfully converged (see Methods).

We found that windows with significantly positive Tajima’s *D* values were older than windows with significantly negative Tajima’s *D* values and windows non significant Tajima’s *D* values (F_1,105_ = 5.58, P = 0.0049; Fig. 4). Although host-associated SNPs were, on average, older than windows that did not contain host associated SNPs, the trend was not significant (F_1,105_ = 0.2, P=0.817). Among windows containing host-associated SNPs identified using samples from all sites, the mean TMRCA of windows with significantly positive Tajima’s *D* values were not significantly greater than either windows with significant negative Tajima’s *D* or non-significant Tajima’s *D* values (F_1,74_=2.19, P=0.14).

**Figure 4:**
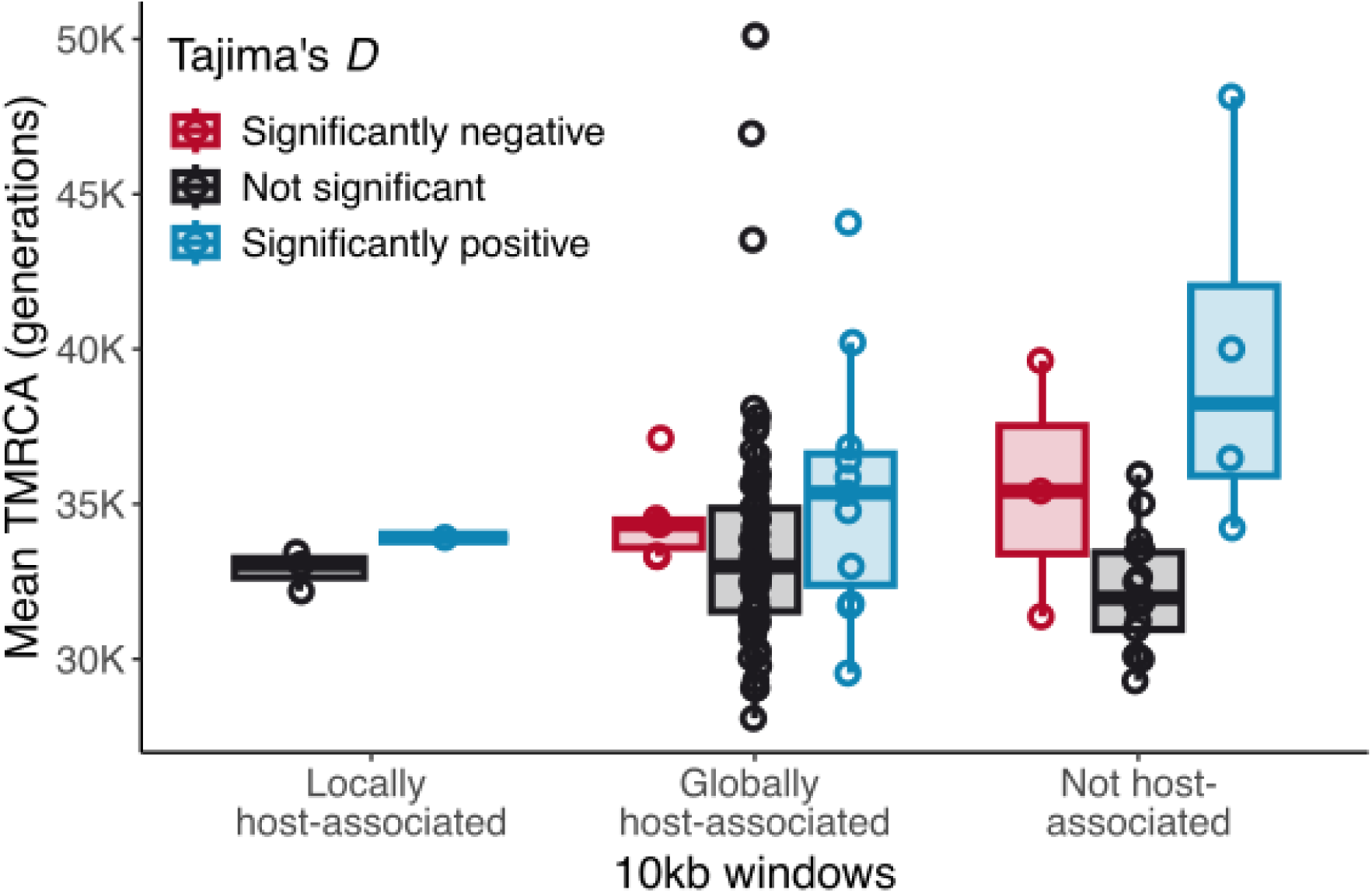
Estimates of time to the most recent common ancestor (TMRCA) of genomic positions containing SNPs. Each point represents the mean TMRCA for all positions within a 1Okb window. Red points and box plots represent 10kb windows with significantly negative Tajima’s D values (treated as evidence of positive selection), black points and box plots represent lOkb windows with non­significant Tajima’s D values.

### Host-associated SNPs are enriched for functions putatively involved in plant cell wall modification and enriched in putative regulatory regions

We performed tests for enrichment on gene ontology (GO) terms on molecular function separately on global host-associated SNPs and local PT host-associated SNPs as well as on SNPs in genomic windows with significantly positive or negative Tajima’s *D* values. For the global host-associated SNPs, four significant GO-terms, glycosyltransferase activity (GO:0016757), hexosyltransferase activity (GO:0016758), glucuronosyltransferase activity (GO:0016757), and UDP-glycosyltransferase (GO:0008194) may play important roles in plant cell wall modification (Table 1). We observed a similar pattern of GO-term enrichment for global SNPs in 10kn windows with significantly negative Tajima’s *D* values (Supplementary Table S2). In contrast, global SNPs localized to 10kb windows with significantly positive Tajima’s *D* values are enriched for various GO terms involving acyltransferases, a broad class of enzymes often involved in lipid metabolism and membrane biosynthesis (Supplementary Table S3).

**Table 1:** Significantly enriched GO-terms for globally host-associated SNPs.

| GO ID | Term | Annotated | Significant | Expected | P-value |
| --- | --- | --- | --- | --- | --- |
| GO:0016757 | glycosyltransferase activity | 51 | 9 | 2.58 | 0.00083 |
| GO:0016758 | hexosyltransferase activity | 46 | 8 | 2.33 | 0.00179 |
| GO:0015020 | glucuronosyltransferase activity | 28 | 6 | 1.42 | 0.00226 |
| GO:0004066 | asparagine synthase (glutamine-hydrolyzing) activity | 2 | 2 | 0.1 | 0.00254 |
| GO:0004703 | G protein-coupled receptor kinase activity | 2 | 2 | 0.1 | 0.00254 |
| GO:0008194 | UDP-glycosyltransferase activity | 32 | 6 | 1.62 | 0.00459 |
| GO:0003824 | catalytic activity | 1073 | 68 | 54.3 | 0.01036 |
| GO:0016884 | carbon-nitrogen ligase activity, with glutamine as amino-N-donor | 4 | 2 | 0.2 | 0.01426 |
| GO:0004674 | protein serine/threonine kinase activity | 57 | 7 | 2.88 | 0.02317 |
| GO:0008237 | metallopeptidase activity | 58 | 7 | 2.93 | 0.02527 |
| GO:0016879 | ligase activity, forming carbon-nitrogen bonds | 14 | 3 | 0.71 | 0.03062 |
| GO:0004222 | metalloendopeptidase activity | 48 | 6 | 2.43 | 0.03206 |
| GO:0016740 | transferase activity | 369 | 26 | 18.67 | 0.04535 |

Host-associated SNPs identified locally at the PT site were mostly enriched for GO-terms with general molecular functions, such as ion binding (GO:0043167), nucleotide binding (GO:0000166), and transmembrane transport activity (GO:0022857) (Table 2). local host-associated SNPs in regions with significantly positive Tajima’s *D* values were enriched for various symporter activities, but these tests are likely underpowered (Supplementary Table S4). No local host-associated SNPs were found in regions with significantly positive Tajima’s *D* values.

**Table 2:** Significantly enriched GO-terms for locally host-associated SNPs.

| GO ID | Term | Annotated | Significant | Expected | P-value |
| --- | --- | --- | --- | --- | --- |
| GO:0043167 | ion binding | 188 | 9 | 2.69 | 0.00053 |
| GO:0005524 | ATP binding | 74 | 5 | 1.06 | 0.00303 |
| GO:0030554 | adenyl nucleotide binding | 75 | 5 | 1.07 | 0.00322 |
| GO:0032559 | adenyl ribonucleotide binding | 75 | 5 | 1.07 | 0.00322 |
| GO:0035639 | purine ribonucleoside triphosphate binding | 79 | 5 | 1.13 | 0.00405 |
| GO:0017076 | purine nucleotide binding | 80 | 5 | 1.15 | 0.00428 |
| GO:0032555 | purine ribonucleotide binding | 80 | 5 | 1.15 | 0.00428 |
| GO:0043168 | anion binding | 80 | 5 | 1.15 | 0.00428 |
| GO:0032553 | ribonucleotide binding | 81 | 5 | 1.16 | 0.00452 |
| GO:0097367 | carbohydrate derivative binding | 81 | 5 | 1.16 | 0.00452 |
| GO:0005509 | calcium ion binding | 26 | 3 | 0.37 | 0.0053 |
| GO:0000166 | nucleotide binding | 102 | 5 | 1.46 | 0.01209 |
| GO:1901265 | nucleoside phosphate binding | 102 | 5 | 1.46 | 0.01209 |
| GO:0036094 | small molecule binding | 104 | 5 | 1.49 | 0.0131 |
| GO:0005488 | binding | 605 | 14 | 8.67 | 0.01416 |
| GO:0015318 | inorganic molecular entity transmembrane transporter activity | 45 | 3 | 0.64 | 0.02431 |
| GO:0015075 | ion transmembrane transporter activity | 47 | 3 | 0.67 | 0.02728 |
| GO:0005230 | extracellular ligand-gated ion channel activity | 2 | 1 | 0.03 | 0.02846 |
| GO:0097159 | organic cyclic compound binding | 406 | 10 | 5.82 | 0.0381 |
| GO:1901363 | heterocyclic compound binding | 406 | 10 | 5.82 | 0.0381 |
| GO:0022857 | transmembrane transporter activity | 57 | 3 | 0.82 | 0.04492 |

Furthermore, a sequence ontology analysis revealed that global host-associated SNPs were significantly enriched in genic regions, a pattern that appears to be mostly driven by enrichment of SNPs in introns (Fig. 5A). A similar pattern was observed for global SNPs in genomic windows with significantly positive Tajima’s *D* values (Fig. 5C). Global SNPs in genomic windows with significantly negative Tajima’s *D* values were under-enriched in exons and for synonymous and nonsynonymous changes (Fig. 5E). In contrast to global host-associated SNPs, locally host-associated SNPs were under-enriched in genic regions, a pattern that seems to be driven by under enrichment of SNPs in introns (Fig. 5B). Locally host-associated SNPs were also enriched for upstream regions (Fig. 5B).This was also true for locally host-associated SNPs in regions with significantly positive Tajima’s *D* (Fig. 5D).

**Figure 5:**
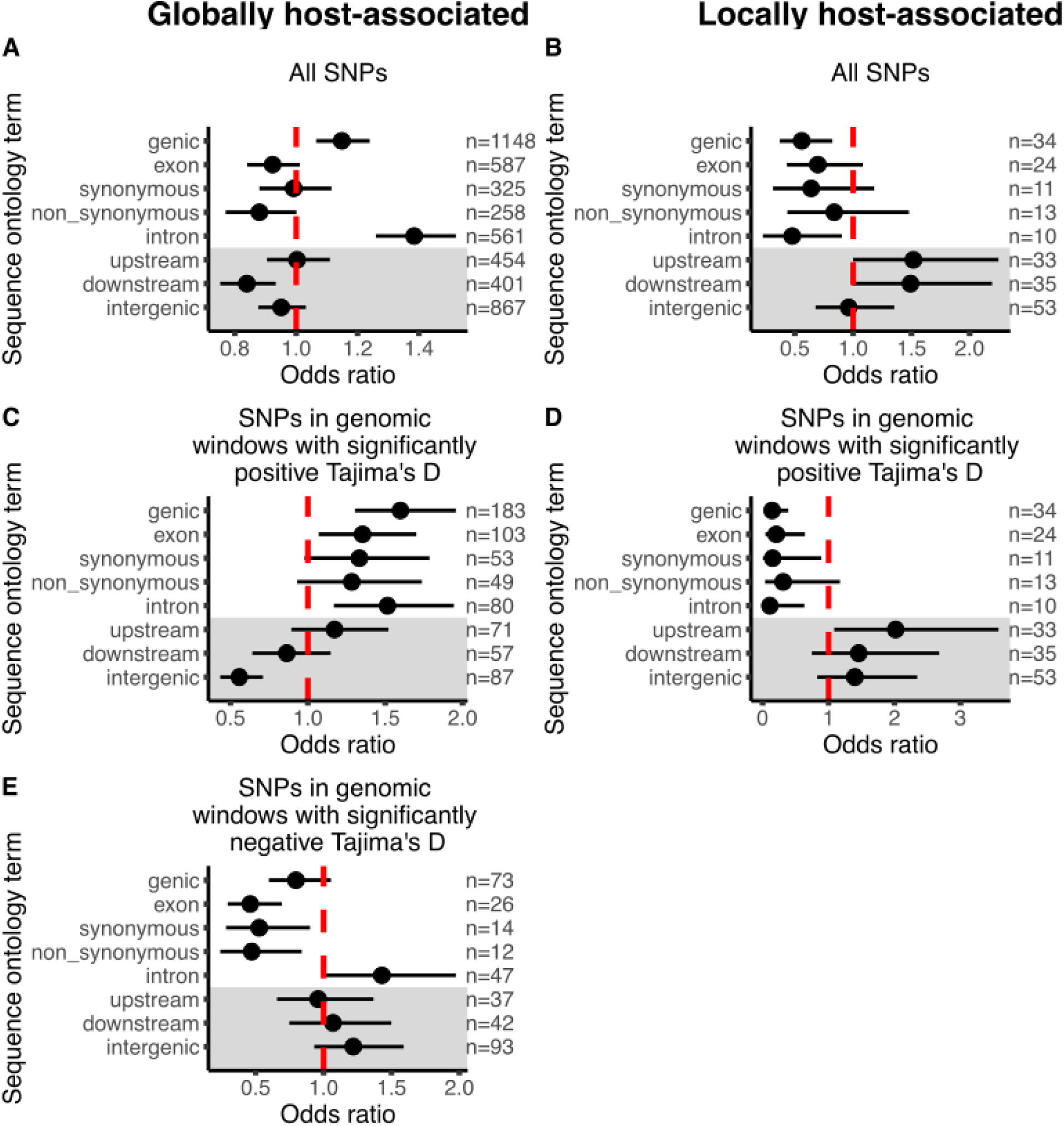
Sequence ontology (SO) enrichment analysis for (A,C,E) global SNPs and (B,D) local PT SNPs. Panels A and B display results of SO enrichment tests on all SNPs from each sample set, panels C and D display results of SO enrichment tests on SNPs residing in 1Okb windows with significantly positive Tajima’s D values, and panel E displays the results of SO enrichment tests on SNPs residing in 10kb windows with significantly negative Tajima’s D values. No local PT SNPs were contained in windows with significantly negative Tajima’s D values. Points are the odds ratios derived from fisher tests and bars are the 95% confidence intervals around the odds ratio. SO terms with error bars that do not overlap with one (the red dashed line) are significantly enriched. SO terms with odds ratios below one (to the left of the red dashed line) are under-enriched, while SO terms with odds ratios above one (to the right of the red dashed line) are enriched. The number of significant SNPs that fall into a given SO term are listed to the right of each plot.

### Evidence for selection on genes and loci with previously identified roles in plant infection

Cross-species eQTL hotspots are regions of the *M. hapla* genome associated with differential expression of hundreds of host plant genes (Guo et al. 2017). We found that global host-associated SNPs were strongly enriched in the four previously identified cross-species eQTL (Fig. 6A). The average Tajima’s *D* of 10kb windows in cross-species eQTL hotspots was higher than the genome-wide average, but not significantly so (F_1,2905_ = 3.323, P=0.07; Fig. 6B).

**Figure 6:**
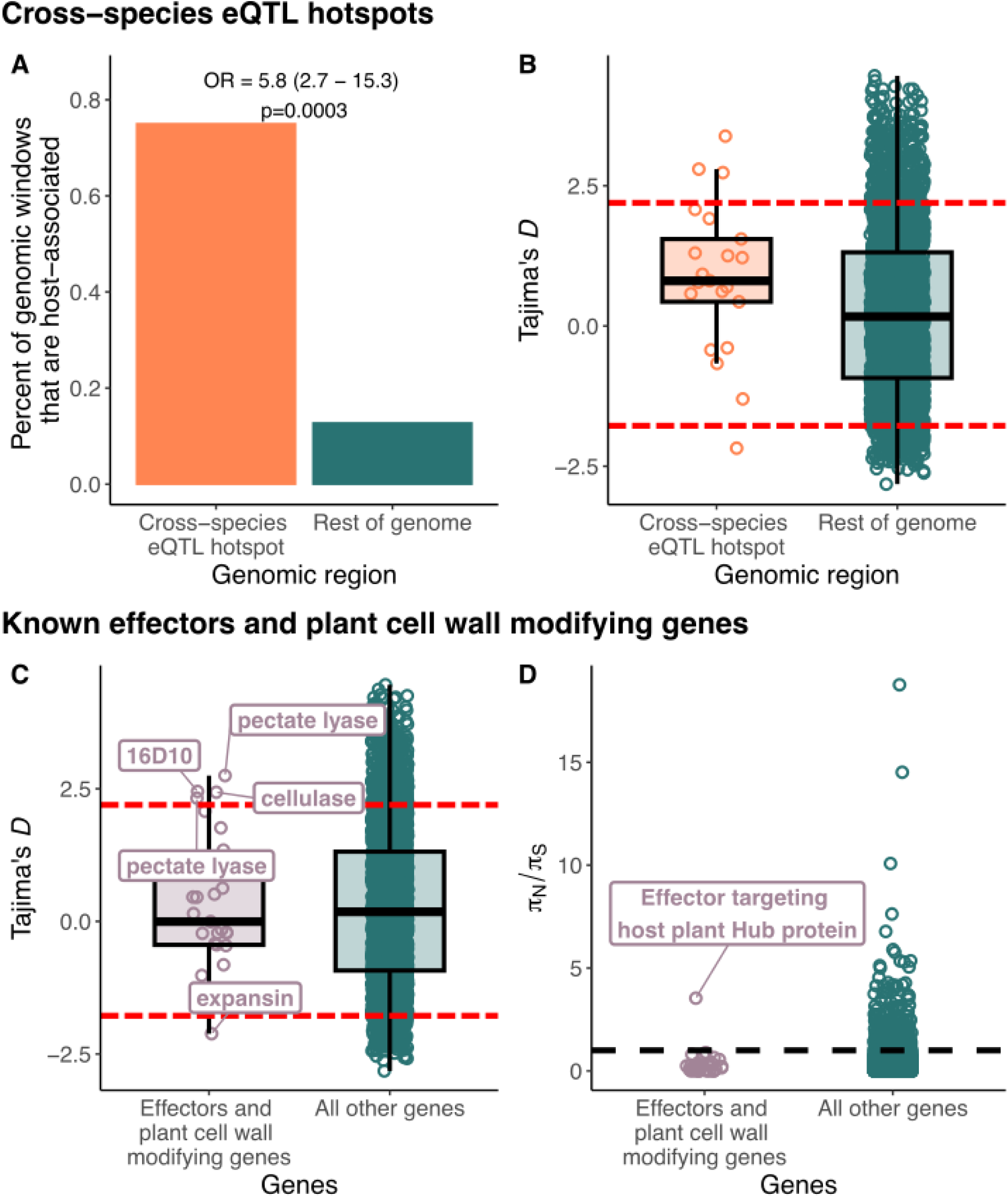
Host-associated variation and signals of selection in loci with known roles in plant infection. (A) Enrichment of 1Okb genomic windows containing globally host-associated SNPs in cross-species eQTL hotspots. (B) Comparison of the distribution of Tajima’s D values for 10kb genomic windows in cross-species eQTL hotspots versus the rest of the genome. (C) Comparison of the distribution of Tajima’s D values for 10kb genomic windows that contain known effectors or plant cell wall degrading enzymes versus the rest of the genome. (D) Comparison of nN/nS estimates for known effectors and plant cell wall degrading enzymes compared to the rest of the genome.

We also searched for host-associated SNPs in a list of 65 effectors and plant cell wall modifying genes that have a functionally validated role in plant infection in *Meloidogyne* (Supplementary Table S5). We found host-associated SNPs in two of these genes, *MhA1_Contig1882.frz3.gene7*, a homolog of the *Msp4* effectors that manipulate host defense response during infection in *M. incognita* and *M. arenaria* (Huang et al. 2003; Przybylska et al. 2023), and *MhA1_Contig1354.frz3.gene2*, a homolog of the *Mj-NULG1a* effector that affects parasite virulence and interacts with plant hub proteins in *M. javonica* (Godinho Mendes et al. 2021). We also found a host-associated SNP in an annotated pectinesterase (*MhA1_Contig555.frz3.gene19*). Pectinesterases are enzymes involved in plant cell wall modification and only exist in the genomes of plants and pathogenic bacteria, fungi, and nematodes.

Five of the known effectors and plant cell wall modifying genes were located in 10kb windows with signals of selection. Specifically, the widely studied effector *Mi16D10* (*MhA1_Contig217.frz3.gene16*) that is required for host infection (Huang, Dong, et al. 2006; Huang, Allen, et al. 2006; Dinh et al. 2015), two pectate lyase genes (*MhA1_Contig2076.frz3.gene1*, *MhA1_Contig331.frz3.gene6*), and a cellulase gene MhA1_Contig802.frz3.gene9) were all located in regions with significantly positive Tajima’s *D* values. An expansin (*MhA1_Contig1277.frz3.gene13*), an enzyme involved in altering the elasticity of plant cell walls, was located in a 10kb window with a significantly negative Tajima’s *D* value. However, as a group, these 65 known effectors and plant cell wall modifying genes did not have Tajima’s *D* values that were significantly different from the rest of the genome (F_1,2905_=0.0011, P=0.97; Fig. 6C).

Finally, we examined signals of selection on the coding sequences of these 65 genes using π_N_/π_S_ (Nei and Gojobori 1986; Hughes and Nei 1988). 38 of the 65 known effectors and plant cell wall modifying genes harbored polymorphism in coding sequences, and all but one had π_N_/π_S_ estimates < 1, indicative of purifying selection (Fig. 6D). There was no difference in the mean π_N_/π_S_ for known effectors and plant cell wall modifying genes compared the rest of the genome (χ^2^ = 0.657, P=0.41). *MhA1_Contig252.frz3.gene17*, which is an ortholog to *Minc00344,* an *M. incognita* gene that interacts with *Glycine max* (soybean) hub proteins and contributes to infection outcomes (Godinho Mendes et al. 2021). has an π_N_/π_S_ estimate of 3.54.

## Discussion

We conducted a population-based resequencing study of root-knot nematode parasites collected from three different host species at multiple geographic sites. Despite no evidence for host-specialized ecotypes, we did find hundreds of loci that appear to be under divergent selection by different host plant species. Many of these SNPs have likely roles in plant infection, including the modification of plant cell walls - a key process in the formation of the root galls that serve as nematode feeding sites (Mitchum and Liu 2022; Bali and Gleason 2024). We also found that a large number of host-associated SNPs are the targets of balancing selection and that regions under balancing selection were, on average, older than the rest of the genome. Furthermore, we found several previously identified effectors appear to also be targets of balancing selection imposed by different hosts. Our results suggest that different host species select for and maintain adaptive diversity in this generalist parasite. This hypothesis represents an important alternative hypothesis to the prevailing co-evolutionary framework in parasite evolutionary genomics: that balancing selection in parasite genomes is indicative of ongoing Red Queen coevolution with a single host. Our study suggests that selection imposed by entire host communities, rather than single taxa, may shape adaptive variation in parasites.

### Host use drives allele frequency differentiation, but not the formation of host-associated ecotypes

Our population structure analyses show that *M. hapla* is a true generalist, rather than a collection of host-specialized ecotypes. (Fig. 1). This is consistent with what is known about *M. hapla* life history: root knot nematodes, including *M. hapla,* are weak dispersers, and thus have limited control over which hosts are available for them to infect (Prot 1980; Pinkerton et al. 1987). Such conditions are predicted to favor the evolution of true generalists (Levins 1968; Felsenstein 1976; Kassen 2002). Our finding that *M. hapla* is a true generalist may also explain its status as a common and destructive agricultural pest. Spillovers and cross-species transmissions are expected to be far more frequent in true generalists (Woolhouse et al. 2001; Power and Mitchell 2004).

Despite *M. hapla’*s identity as a true generalist, we found several hundred SNPs with differentiated allele frequencies between nematodes collected from different host plants indicating that *M. hapla* populations do respond to differential selection by hosts (Fig. 2).Contrary to our initial expectation, the comparison between nematodes isolated from the most closely related host pair—*M. lupulina* and *T. repens*—produced some of the strongest patterns of allele frequency differentiation in our dataset. This finding suggests that, at least in some cases, the response to differential host selection is independent of host phylogeny. Weak phylogenetic signal in parasite adaptation to different hosts is possible if phylogenetically distant hosts share traits that produce similar selective pressures (Futuyma and Moreno 1988; de Vienne et al. 2013). For example, both *T. repens* and *L. vulgare* have shallow and non-woody root systems relative to *M. lupulina* (Chmelíková and Hejcman 2012). Root physiology is likely important in *M. hapla* infection success (Eisenback and Triantaphyllou 1991).

Consistent with root physiology shaping selection, we found that globally host-associated SNPs were enriched for genes with likely functions in plant cell wall modification. Specifically, globally host-associated SNPs were overrepresented in genes involved in glycosyltransferase, hexosyltrasferase, and glucuronosyltransferase activity; all enzymes that, in plants, are essential for synthesizing plant cell wall components like cellulose (Amos and Mohnen 2019).

We also found host-associated SNPs in a pectinesterase (*MhA1_Contig555.frz3.gene19*), which is involved in the degradation of pectin, another key plant cell wall component (Opperman et al. 2008; Danchin et al. 2010). We hypothesize that these genes in *M. hapla* facilitate the modification of plant cells into the giant cells that serve as the root knot nematode feeding site. Additionally, the observation of host-associated SNPs in genes that manipulate the plant immune response suggest that plant cell wall modification is likely not the sole target of selection.

We also found that host-associated SNPs were overrepresented in regulatory regions. Global host-associated SNPs were enriched in introns, which implicates a potential role for variation in the expression of specific gene isoforms, via changes to splice sites or splicing regulatory sequences, underpin the response to differential host selection. In comparison, local host-associated SNPs were enriched upstream of start codons, a region that may contain promoter and enhancer sequences or to transcription factor binding sites. Enrichment of host-associated SNPs in putative regulatory regions is consistent with the expectation that regulatory mutations are under weaker selective constraint than coding mutations (Wray 2007; Barrett and Schluter 2008).

To summarize, we found hundreds of SNPs that are putative targets of differential host selection in this generalist parasite. This finding complements and extends theory on the evolution of generalist parasites in two ways. First, the maintenance of polymorphisms in a true generalist is consistent with theory and previous studies on evolution in heterogeneous environments (Levins 1968; Felsenstein 1976; Kassen 2002). Second, the apparent variation in selection exerted by different host species highlights the important role that host community composition plays in the evolution of generalist parasites (Woolhouse et al. 2001; Power and Mitchell 2004; Thompson 2005). Our finding of weak genetic differentiation between nematodes isolated from distantly related host plants (*L. vulgare* and *T. repens)* raises the possibility that the phylogenetic structure of host communities may be less important in the evolution of generalists than specialized parasites (Parker et al. 2015).

### Long-term balancing selection maintained by heterogeneous selection across host species

One of the most striking patterns in our dataset is the enrichment of host-associated loci in genomic windows exhibiting a signal of balancing selection as measured by Tajima’s *D* (Fig. 3). A similar trend also exists when comparing the average Tajima’s *D* of windows containing host-associated SNPs versus those that do not. Our confidence that these represent signals of balancing deriving from differences in selection imposed by different hosts is strengthened by our observations that these regions tend to be, on average, older than the rest of the genome (Fig. 4). Together, our data suggest that heterogenous host selective pressures maintain balanced polymorphisms at loci involved in host infection.

An alternative explanation for the long-term maintenance of host-associated polymorphisms is specific coevolution within one or several different host species (Thompson 1994). However, given the high variability in which root knot nematodes encounter different host species, it is unlikely that they infect any specific host species consistently enough to allow coevolution to occur (Gomulkiewicz et al. 2000; Gibson 2019).

Three genes involved in cell wall modification and one involved in host plant developmental reprogramming exhibited signals of balancing selection, and one gene, also involved in plant cell modification, exhibited a signal of positive selection. One of the genes under balancing selection, *16D10* (*MhA1_Contig217.frz3.gene16*) facilitates gall formation by manipulating *CLAVATA3/EMBRYO SURROUNDING REGION (CLE)* signaling in host plants (Nakagami et al. 2023). The one gene under positive selection is an expansin (MhA1_Contig1277.frz3.gene13) involved in altering the elasticity of plant cell walls. The fact that it is under positive selection suggests that itis globally beneficial across all host plant species in our study.

Finally, our data suggest that the coding sequences of known effectors are subject to strong selective constraint. All 65 known effectors and plant cell wall modifying genes had π_N_/π_S_ < 1, consistent with purifying selection. The one exception was *MhA1_Contig252.frz3.gene17*, a homolog of the *M. incognita* gene *Minc00344,* which binds to host plant hub genes and influence nematode infection success (Godinho Mendes et al. 2021). Generally π_N_/π_S_ > 1 are interpreted as signals of multiallelic balancing selection, as the number of nonsynonymous mutations that need to be simultaneously segregating to detect π_N_/π_S_ values above one is inconsistent with positive selection (Hahn 2019). Several immune-related genes in other systems, such as the major histocompatibility complex genes, also contain high values of π_N_/π_S_ presumably because diffuse interactions with several parasite taxa works to maintain many distinct alleles of these genes (Hughes and Nei 1988; Spurgin and Richardson 2010; Osborne et al. 2017; Radwan et al. 2020).

### Evolutionary genomics of wild parasites

We present evidence that infection-associated SNPs are maintained by balancing selection resulting from diffuse interactions with different host species. The idea that diffuse interactions among multiple parasites or hosts can support the long-term maintenance of polymorphisms is not new, but has been largely explored from the perspective of hosts interacting with several parasite species (Karasov et al. 2014; Radwan et al. 2020). Given that the majority of parasite species are capable of infecting more than one host species (Barrett et al. 2009), the maintenance of polymorphism via diffuse selection from different host species is probably not unique to *M. hapla*. Therefore, future research should be careful when ascribing balancing selection in generalist parasites to pairwise host-parasite coevolution.

Parasites are small and can be difficult to isolate, which has led to a lack of whole genome studies, at least relative to host species. The whole genome amplification approach we employed here gave us the opportunity to explore genome-wide patterns of variation which revealed extensive evidence of host-driven selection. This method could also be applied in a laboratory setting to quantify the fitness effects of alternative alleles on different hosts to interrogate putative genetic tradeoffs identified in population genomic studies of selection like ours. Whole genome amplification could be an invaluable tool to study the evolutionary dynamics of wild parasites (Shortt et al. 2017; Cruaud et al. 2018; O’Grady et al. 2022; Pilling et al. 2023).

## Methods

### Sample collection, whole genome amplification, and sequencing

Infected plants belonging to three host plant species (*Medicago lupulina*, *Trifolium repens*, and *Leucanthemum vulgare*) were collected at each of four study locations, three in Virginia (Boley Field Road (BR), Smallville Fire Access Road (SM), and Pott’s Mountain Trail (PT)), and one in Pennsylvania (State Game Lands 39 (PA)) (Fig. 1A). After field collections, galls were cut from roots and stored at −80 °C.

Whole nematodes were dissected from frozen gall samples and again stored at −80 °C. Next, we performed whole-genome amplification on individual whole nematodes (See supplementary methods for full protocol). We confirmed species identity of amplified samples with PCR using the diagnostic primer set JMV1 and JMVhapla reported in Wishart et al. (2002). The amplified DNA was then sent to Admera Health Inc. (South Plainfield, NJ, USA) for library preparation using the KAPA HyperPrep PCR-free Kit (Roche Diagnostics Corp., Indianapolis, IN, USA) and sequencing on an Illumina NovaSeq machine. We targeted 30x coverage (∼1.56 Gb) for each sample.

### Mapping and Quality Control

After verifying that all samples passed quality control with fastQC (Andrews 2010) and trimming Illumina adaptor sequencing with trimmomatic v.0.36 (Bolger et al. 2014), we aligned raw reads against the *Meloidogye hapla* reference genome available on Wormbase ParaSite v.18 (Opperman et al. 2008; Howe et al. 2016; Howe et al. 2017) using the BWA-MEM algorithm implemented in BWA v.0.717 (Li and Durbin 2009). We used Picard-tools v.1.141 (Anon 2019) to remove duplicated reads.

Whole genome amplification relies on large sets of random hexamer primers, which can potentially result in uneven coverage (O’Grady et al. 2022). We estimated coverage breadth, average coverage depth, and coverage evenness for each sample (Oexle 2016)(Supplementary table S6). We also computed pairwise correlation coefficients for read counts in 1-kb bins between all individuals to ensure that the expected uneven read distribution resulting from WGA was consistent across samples (Supplementary Figure S4).

### Variant calling and filtering

We called variants on the 70 individuals that had at least 80% of reads with a primary alignment to the reference genome (Supplementary table S6). We followed the GATK v4.1.7.0 best practices for germline variant calling, which involves using GATK’s HaplotypeCaller and genotypeGCVF algorithms (Supplemental methods). We further filtered our variant set by removing indels, retaining only biallelic sites, removing variants in low complexity regions, removing variants with less more than 80% missigness, removing variants with a minor allele counts of less than 3, and removing variants that deviated from Hardy-Weinberg equilibrium. After all filtering we retained a list of 117,218 biallelic SNPs.

### Population structure

We inferred population structure using two complementary methods: a standard PCA and admixture analysis. Both analyses require sets of unlinked SNPs, so we pruned SNPs with pairwise LD values above 0.2 using PLINK v2.0 in sliding windows containing 50 SNPs and advancing 5 SNPs at a time (Chang et al. 2015; Purcell and Chang). PCA was performed using the PCA command in the R package SNPrelate (Zheng et al. 2012). Admixture analysis was performed using fastStructure (Raj et al. 2014), implemented in the wrapper program structure_threader (Pina-Martins et al. 2017). We ran fastStructure using admixture proportions of *K*=1 to *K*=15. We used the Evanno method to select the best fitting admixture proportion *K* (Evanno et al. 2005).

### Identification of host-associated SNPs

We used two complementary approaches to identify host-associated SNPs using the program BayPass (Gautier 2015; Olazcuaga et al. 2020). The first approach leveraged all 59 individuals collected from the three sampling sites (BR, SM, and PT) in Virginia (“global”) and the second used only the 29 individuals collected from our most well sampled site at PT (“local”). For both approaches, we performed three Contrast Analyses (between nematodes collected from each pair of host species) implemented in BayPass. Contrast analyses are a categorical variant of genotype-environment association (GEA) model. Full details of how BayPass was implemented appear in the supplementary materials. Briefly, both the “core” and “contrast” models were run using default priors for 25,000 MCMC interactions with a 5,000 iteration burn-in. The contrast model was provided with the the Ω matrix estimated from the core model so that population stratification could be controlled for during estimation of the *C_2_* statistics. Three independent model runs resulted in highly correlated *C_2_* indicating model convergence.

We calibrated *C_2_* estimates using a pseudo-observed dataset (POD) (Supplementary methods; Gautier 2015). We classified SNPs with *C_2_* estimate greater than the 0.99 percentile of the POD as host-associated.

### Inference of selection on host-associated loci

To infer selection on host-associated SNPs, we quantified Tajima’s *D* in non-overlapping sliding windows of 10kb using the –TajimaD command in vcftools (Danecek et al. 2011). 10kb roughly equates to the genome-wide average recombination distance of 143 cM/Mb (Thomas et al. 2012). Because our dataset contains evidence of strong population structure, distant population split times, and limited migration among populations (see Results) we performed our selection inference on our most well-sampled study site, PT, rather than the combined dataset (Simonsen et al. 1995).

We established thresholds for detecting significant balancing and positive selection using the 0.5% and 95.5% quantiles of a distribution of Tajima’s *D* estimated using simulated genetic data (see ‘Demographic history estimation and genetic simulations using fastsimcoal’ section). We used Fisher’s exact tests to assess whether 10kb genomic windows that contained host-associated SNPs were enriched for significantly positive or negative Tajima’s *D* estimates. We also used a simple linear model to test whether the average Tajima’s *D* estimate of 10kb genomic windows that contain host-associated SNPs were different from those that do not contain host-associated SNPs.

### Ancestral recombination graph (ARG) estimation

To estimate the age in generations of alleles under balancing selection, we inferred ancestral recombination graphs (ARG) for the PT population sample on the approximately 250 contigs of the *M. hapla* assembly that contained host-associated SNPs using ARGweaver (Rasmussen et al. 2014). We followed Hubisz & Siepel (2020) to run ARGweaver (Supplementary methods).

### Sequence and gene ontology enrichment analysis

To test whether host-associated SNPs were enriched in specific sequence features we performed an enrichment analysis using sequence ontology assignments made with snpEff (Cingolani et al. 2012). We first constructed a snpEff genome data for *M. hapla* with the assembly in the WormBase v.18 release. We then queried our full set of 117,218 filtered SNPs against this database to get predicted effect annotations for each SNP. Next, we used the resulting predicted annotations to perform Fisher exact tests to determine whether host-associated SNPs were enriched in the sequence features presented in Figure 5. Genic regions were defined as all exon and intron positions, upstream regions were defined as all positions 1kb upstream of the first start codon, and downstream regions were defined as all positions 1kb downstream of the final stop codon.

To test whether host-associated SNPs are enriched for specific molecular functions, we performed a gene ontology (GO) term enrichment analysis using the R package topGO (Alexa and Rahnenfuhrer). GO term enrichment was performed using the 9,429 genes in the *M. hapla* assembly that have annotated GO terms. To test for significant enrichment we used the “classic” algorithm and performed Fisher exact tests.

### Demographic history estimation and genetic simulations using fastsimcoal

To establish significance thresholds for selection inference using Tajima’s *D*, we simulated genetic data from a best-fitting demographic model in fastsimcoal v.2.7 (Excoffier et al. 2021). Detailed methodology for demographic inference and genetic simulations are reported in the supplemental methods. Briefly, We tested four different historical migration scenarios: strict isolation (SI, i.e. no migration among demes), continuous migration among all demes for the entire demographic history, continuous migration before the second deme fission only, and continuous migration after the second deme fission only. We tested these four different scenarios on four different tree topologies: three different bifurcating trees and one multifurcating tree, for a total of 16 models.

To establish significance thresholds for selection inference with Tajima’s *D*, we performed 100 independent genetic simulations according to the best fitting demographic model using fastsimcoal. For each simulation, initial parameter values were randomly drawn from the 90% bootstrap confidence interval. We quantified Tajima’s *D* in the resulting simulated genetic data in non-overlapping sliding windows of 10kb using the –TajimaD command in vcftools. The bottom 0.5% and top 99.5% quantile of the distribution of Tajima’s *D* values were used as the threshold for detecting significant positive and balancing selection, respectively (Supplementary Methods).

## Author Contributions

M.C. and C.W. conceived of the study and designed the sampling scheme. M.C., A.C., A.G., and L.W. collected the samples. M.C. performed data analysis with continual feedback from all authors. M.C. wrote the initial draft of the manuscript and all authors contributed to revisions.

## Acknowledgments

We are grateful to the United States Forest Service, the Pennsylvania Game and Fish Commission, The University of Virginia Mountain Lake Biological Station, and the University of Pittsburgh Pymatuning Lab of Ecology for granting us permission to sample root knot nematodes on their land. We thank Lauren Kerwien, Emile Gluck-Thaler, Samantha Catella, and Dana Crawford for assistance with field sampling, Wil Prall for help with developing our whole genome amplification protocol, John Wagner for allowing us to use his lab’s fluorometer, and Junhyong Kim, Erol Akçay, Scott Poethig, Paul Schmidt, and Kattie Barrot for helpful discussions. We also thank Addison Buxton-Martin, Chang-Yu Chang, Anisa Robinson, Eunnuri Yi, and Jazmine Rud for providing comments on this manuscript. This work was funded by NSF-DEB 2118397 to Corlett Wood.

## Data Accessibility Statement

Raw sequence reads will be deposited at NCBI SRA. The VCF containing the variants on which analyses were performed along with all analysis scripts will be deposited on Dryad.

