## Supplementary Methods for "Evidence for balancing selection on loci associated with host use in the genome of a generalist plant parasite"

*Whole genome amplification of whole nematodes*

All of the reagents mentioned here were contained in the Qiagen REPLI-g Single Cell Kit (cat. # 150343). Prior to performing whole genome amplification, Buffer D2 was prepared using 1 part 1 M DTT and 11 parts Buffer DLB. A 1:10 dilution of 1 M DTT in $H_2O$ sc was also prepared beforehand.

To reduce the chance of inadvertently amplifying bacteria and plant DNA, individual nematodes were washed five times in sterile 1X PBS by placing a row of five 1X PBS droplets on a sterile petri dish and carefully and repeatedly drawing the nematode up and down across the 5 droplets. At the final droplet, individual nematodes were crushed with the end of the pipette tip and were carefully drawn up and down with the pipette many times to homogenize the sample. 3 μL of D2 buffer was added to each homogenized whole nematode sample and mixed carefully by flicking. Samples were then incubated at 65 °C for 10 minutes in a dry bath. 3 μL of Stop Solution was then added to each sample. While samples were incubating, a REPLI-g sc polymerase master mix was prepared using a recipe of 6.5 μL $H_2O$ sc, 29 μL REPLI-g Reaction Buffer, 2.5 μL diluted DTT, and 2 μL REPLI-g sc DNA Polymerase for each sample. 40 uL of master mix was added to each sample and samples were incubated at 30 °C for three hours. After this incubation, samples were incubated at 65 °C for 3 minutes to inactivate the REPLI-g sc DNA Polymerase. Amplification products were quality checked using nanodrop and gel electrophoresis. DNA was quantified using a PicoGreen assay. Reactions resulted in approximately 100-600 ng/μL of amplified product for each sample (Supplementary Table S7).

*Phylogenetic analysis of de-novo mitochondria genome assemblies*

*M. hapla* occurs in at least two cytological forms (Eisenback and Triantaphyllou 1991). One form reproduces via facultative meiotic parthenogenesis (i.e. selfing) and the other by obligate mitotic parthenogenesis (i.e. asexual) and they are therefore reproductively isolated. These forms also differ in their chromosome number and ploidy. To determine whether there could be cytological or other subspecific variation in our dataset, we constructed a phylogenetic tree from de-novo mitochondria assemblies. We used GetOrganelle with default settings to assemble mitochondrial genomes for each individual and used the default GetOrganelle animal mitochondrial database to seed the assemblies (Jin et al. 2020). We then used Ska2 to construct a split *k*-mer based alignment and to perform SNP calling on this set of mitochondria assemblies (Harris 2018). Using these SNPs, we constructed a neighbor-joining tree using VeryFastTree (Piñeiro et al. 2020).

*Variant calling and hard filtering*

We followed the GATK v.4.1.7.0 best practices workflow for germline variant calling to call variants in our dataset (McKenna et al. 2010). First we used Picard-tools v.1.141 to generate a reference sequence dictionary required by GATK (Picard Toolkit 2019). Then, we used GATK's HaplotypeCaller function to call SNPs and Indels. HaplotypeCaller is computationally intensive, so we employed an array job whereby an independent instance of HaplotypeCaller was run on each sample. Individual sample genotypes were then combined using GATK's combineGVCF command. Variants were then called on this combined genotype file using GATK's genotypeGCVF command. We filtered variants using the standard hard filtering thresholds recommended by the GATK v.4.1.7.0 best practices workflow. Specifically, we filtered SNPs with a quality depth (DP) of less than 2, a mapping quality (MQ) of less than 40, fisher strand (FS) of greater than 60, a MQ rank sum of less than -12.5, and read position rank sum of less than -8.

*Additional BayPass details*

BayPass requires a table of allele frequency counts for each SNP and population in the dataset, which we generated using the -freq counts command in PLINK 2.0. Because we were interested in identifying SNPs associated with host adaptation, we calculated allele frequency counts for nematodes collected from each host plant in each geographic location for a total of 9 populations. We first ran the "core" BayPass model with just the population allele frequency counts to estimate the scaled variance-covariance matrix **Ω** of population allele frequencies. We ran the core model with default priors, 25,000 MCMC iterations with a burn-in of 5,000 iterations. We ran the core mode 5 times, each with a different seed for the random number generator, to assess model convergence. We found nearly perfect correlations for the posterior means of the elements of **Ω** matrices across model runs (pearson's $r > 0.999$ for all pairwise comparisons. Thus, we used one randomly chosen **Ω** matrix for input in downstream models (Supplementary Figure S5). Next, we ran the "contrast" model in BayPass, which, for each SNP, estimates the mean squared difference of the sum of standardized allele frequencies, called the $C_2$ statistic, of two groups of individuals or populations.

*BayPass POD generation*

The *p*-values of $C_2$ estimates were misbehaved (Supplementary figure S6), potentially owing to the small sample and small number of SNPs used in our study (Gautier 2015; Olazcuaga et al. 2020). To generate a POD, we simulated population allele frequency counts using the empirical **Ω** matrix estimated from the core model. We fed the POD through the 'core' BayPass model and found that the posterior estimate of the POD **Ω** matrix was close to the empirical **Ω** matrix (Forstner and Moonen Distance = 1.086). The similarity of POD and empirical **Ω** matrices indicates that the POD is faithfully mimicking the real dataset. Next, we performed the contrast model for each pairwise comparison of our host-associated groups of nematodes using the simulated allele counts contained in the POD.This produced what is effectively a null distribution of $C_2$ statistics for each of our three host-associated nematode group comparisons that takes into account the shared ancestry of the populations under study.

*Ancestral recombination graph inference*

ARGs are a complete record of all recombination and coalescence events since divergence among sequences in a population sample (Rasmussen et al. 2014). As such, ARGs represent a complete genealogy of every genomic position under study. We used ARGs to derive estimates of the time to the most recent common ancestor (TMRCA) of 10 bp compressed genomic blocks. If a given locus is under balancing selection, then its TMRCA should usually be longer than a locus under weak selective constraints (i.e. neutral).

To estimate ARGs, we first generated mask files for each individual in our sample to mask missing genotype calls. We then used the command `arg-sample` to infer ARGs using a VCF file containing all individuals from PT and all 117,218 SNPs. Separate instances of arg-sample were executed on single contigs at a time. We executed arg-sample with the following parameters: a VCF minimum quality score of 30 (`--vcf-min-qual 30`), masked any windows of 5 bp that had 2 or more SNPs (`--mask-cluster 2,5`), weight genotypes by their phred-likelihoods (`--use-genotype-probs`), a genome-wide mutation rate of 2.0 x $10^{-8}$ (`-m 2.0e-8`), a genome-wide recombination rate of 1.4x$10^{-6}$ (`-r 1.4e-6`), a maximum time point in the model in units of generations of 100 x $10^5$ (`--maxtime 100e5`), set the delta value for choosing log times as 0.01 (`--delta 0.01`), and set the site compression at 10 bp (`-c 10`). All other parameters were left on their default settings. We ran 5000 iterations for each model (`--n 5000`) with the default thinning interval of 10. Owing to the small size of many contigs, many models failed to converge. We only used contigs that had ARG models with sufficient convergence in downstream estimation of TMRCA. We used the command `arg-summarize` to derive mean and standard deviation calculation of TMRCA estimates across MCMC iterations. We used a burnin of between 2000 and 3000 iterations depending on the patterns of convergence for models on different contigs.

$\pi_N/\pi_S$ *Estimation*

To compute $\pi_N/\pi_S$ , we followed the method outlined in Hughes and Nei (1988). Briefly, we first generated consensus sequences in fasta format using bcftools for all 14,419 genes in the *M. hapla* assembly for all 66 individuals in our dataset that had more than 80% of reads mapping to the *M. hapla* assembly. We then concatenated consensus sequences for all individuals to generate individual multiple alignment FASTAs (MAF) for each individual gene. For each gene, we computed $d_N$ and $d_S$ for all pairwise combinations of sequences using the method of Nei and Gojobori (1986) implemented in the BioPython (Cock et al. 2009) function `calculate_dn_ds`. Next, we calculated the mean $d_N$ and mean $d_S$ of all pairwise sequence combinations to estimate $\pi_N$ and $\pi_S$, respectively. This allowed us to generate $\pi_N/\pi_S$ estimates for all genes.

*Demographic history estimation and genetic simulations using fastsimcoal*

We performed our demographic inference in fastsimcoal v.2.7 (Excoffier et al. 2021). Prior to analysis with fastsimcoal, we removed SNPs located in genic regions using bedtools (Quinlan and Hall 2010), and pruned SNPs in high LD (using a cut of < 0.2) in PLINK2.0 (Chang et al. 2015). We then used easySFS (Gutenkunst et al. 2009) to generate a joint three-dimensional minor allele SFS (i.e. a folded SFS) to use as input into fastsimcoal. We used easySFS's preview function to select the optimal projection.

As mentioned in the main text, We tested four different historical migration scenarios: strict isolation (SI, i.e. no migration among demes), continuous migration among all demes for the entire demographic history, continuous migration before the second deme fission only, and continuous migration after the second deme fission only. We tested these four different scenarios on four different tree topologies: three different bifurcating trees and one multifurcating tree, for a total of 16 models. For each demographic model, we fit parameters to the observed joint three dimension minor allele SFS using 200,000 coalescent simulations and 50 ECM parameter optimization cycles. 100 independent fits were performed for each model to randomize starting parameter values to increase the chance that we found the global optimum for the combination of parameter values in each of our demographic models.

To select the best fitting demographic model, we first identified the best run of each of the 16 demographic models based on which had the highest likelihood. Then, for each of the 16 demographic models, we performed 100 model runs using the parameters from the initial run with the highest likelihood. These 100 model runs were used to produce likelihood distributions. A continuous migration model with the BR population as the outgroup had the highest likelihood distribution and was selected as the best model. This demographic model also had the best AIC score.

To assess certainty in demographic model parameter estimates we performed a parametric bootstrapping procedure. This involved first simulating 100 three dimensional folded site frequency spectra using the parameters from the best fitting model. Then, for each independent simulation, we estimated demographic parameters using the same estimation and template files used to identify the original best fitting model. We used the distribution of the estimated parameters from the simulated data to produce a 95% bootstrap confidence interval for each parameter.

To produce significance thresholds for significance testing with Tajima's *D*, we performed 100 independent simulations of genetic data using random draws from the 95% bootstrap confidence intervals for each demographic parameter. For each run, 20 independent chromosomes of 100,000bp in length were simulated. The resulting genotype files were converted to VCFs using custom awk scripts (see dryad repository). Tajima's *D* was estimated in 10kb non-overlapping windows using the –TajimaD command in vcftools, just as was done for the empirical data. The resulting simulated distribution of Tajima's *D* values effectively serves as a null expectation of nucleotide variation in the absence of selection. Furthermore, this approach also allows us to reasonably control for the effect of population stratification, population contraction and expansion, migration, and inbreeding on our inference of selection. The 0.5% and 99.5% quantiles of the distribution of the simulated Tajima's *D* values were used as the significant thresholds for detecting positive and balancing selection, respectively.

### Supplementary Figures and Tables

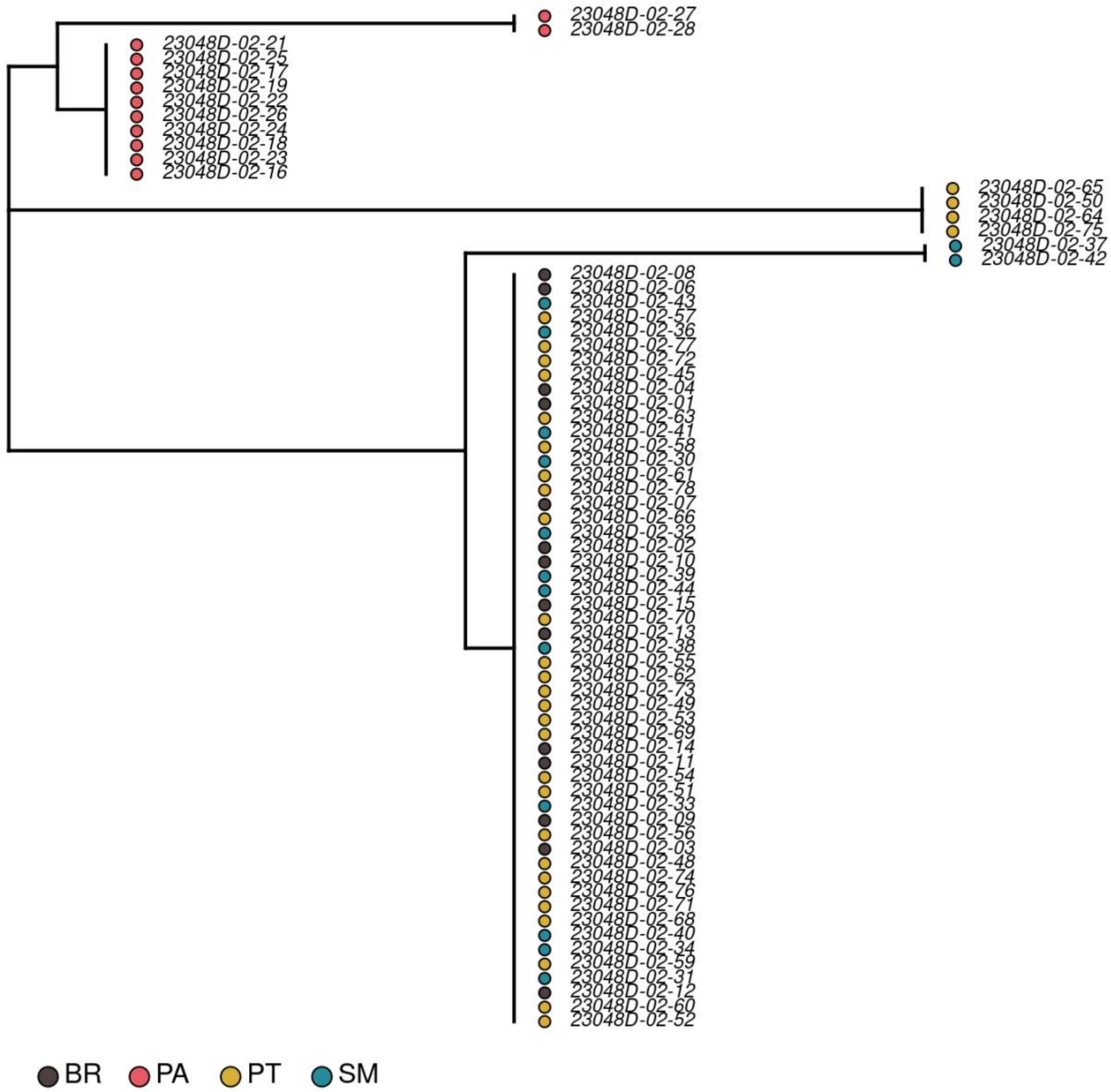


**Supplementary Figure S1**: Neighbor-joining tree of SNP variation in sample mitochondria assemblies. Colors represent sampling location. Three distinct clades were revealed corresponding to all samples from PA, the four individuals from PT with nearly complete assignment to the dominant PA population cluster, and the rest of the samples collected across the three sites in Virginia. Thus, PA individuals and four aforementioned PT individuals may belong to a different cytological type or subspecies of *M. hapla*.

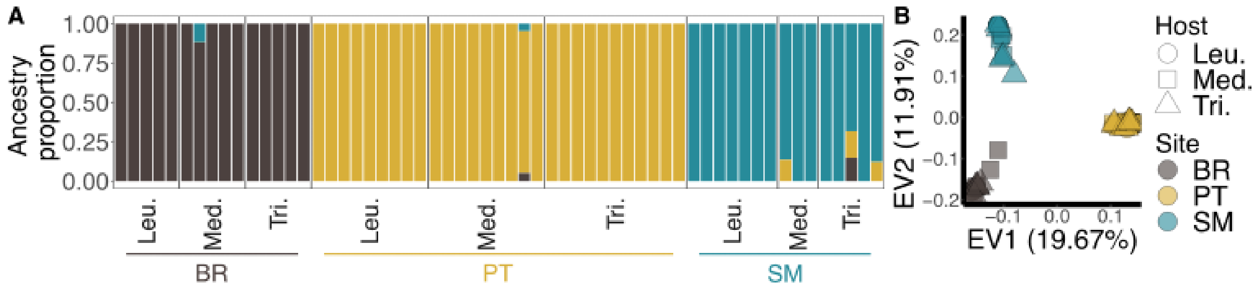


**Supplementary Figure S2**: Population structure of root-knot nematodes collected at sites in Virginia. (A) Ancestry proportions for each individual root-knot nematode calculated using fastSTRUCTURE with *K* = 3 and colored by the dominant cluster at each sampling location. Each bar represents an individual root-knot nematode and are grouped by the host plant species they were sampled from and sampling location. (B) PCA showing the first and second eigenvectors of genetic variation across root-knot nematodes collected at sites near MLBS. Points represent individuals and are colored by sampling location according to the dominant population clusters identified in the fastSTRUCTURE analysis. Point shapes correspond to the host plant species that individuals were sampled from. In each panel, host plant species are abbreviated; Leu. = *Leucanthemum vulgare*, Med. = *Medicago lupulina*, and Tri. = *Trifolium repens*

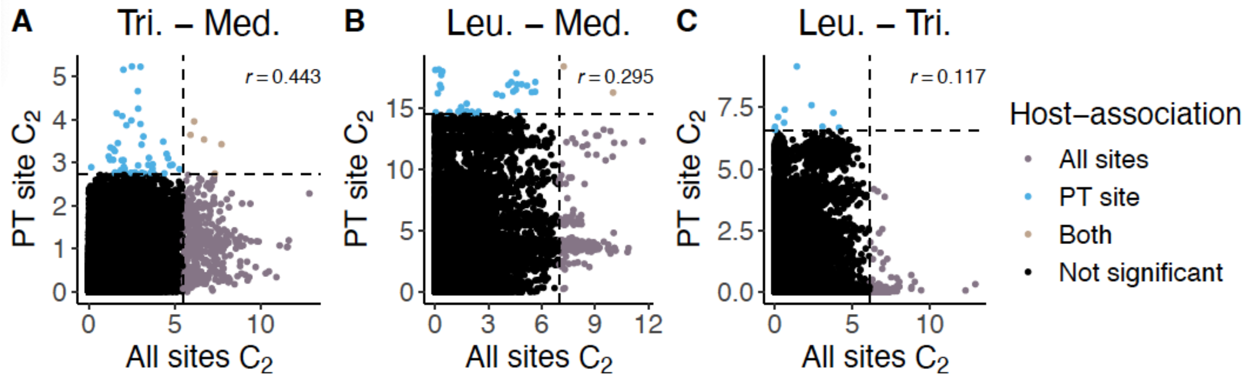


**Supplementary Figure S3**: Plot of $C_2$ estimated leveraging samples from all sites (X-axes) versus $C_2$ estimated using samples within the PT site (y-axes) for comparisons between (A) *T. repens* and *M. lupulina* nematodes (Tri. - Med.), (B) *L. vulgare* and *M. lupulina* nematodes (Leu. - Med.), and (C) *L. vulgare* and *T. repens* nematodes (Leu. - Tri.) . SNPs identified as significantly host-associated using samples from all sites are plotted in purple and SNPs identified as significantly host-associated locally in the PT site are plotted in blue. The dashed lines represent the 1% quantile of the PODs generated for outlier detection.

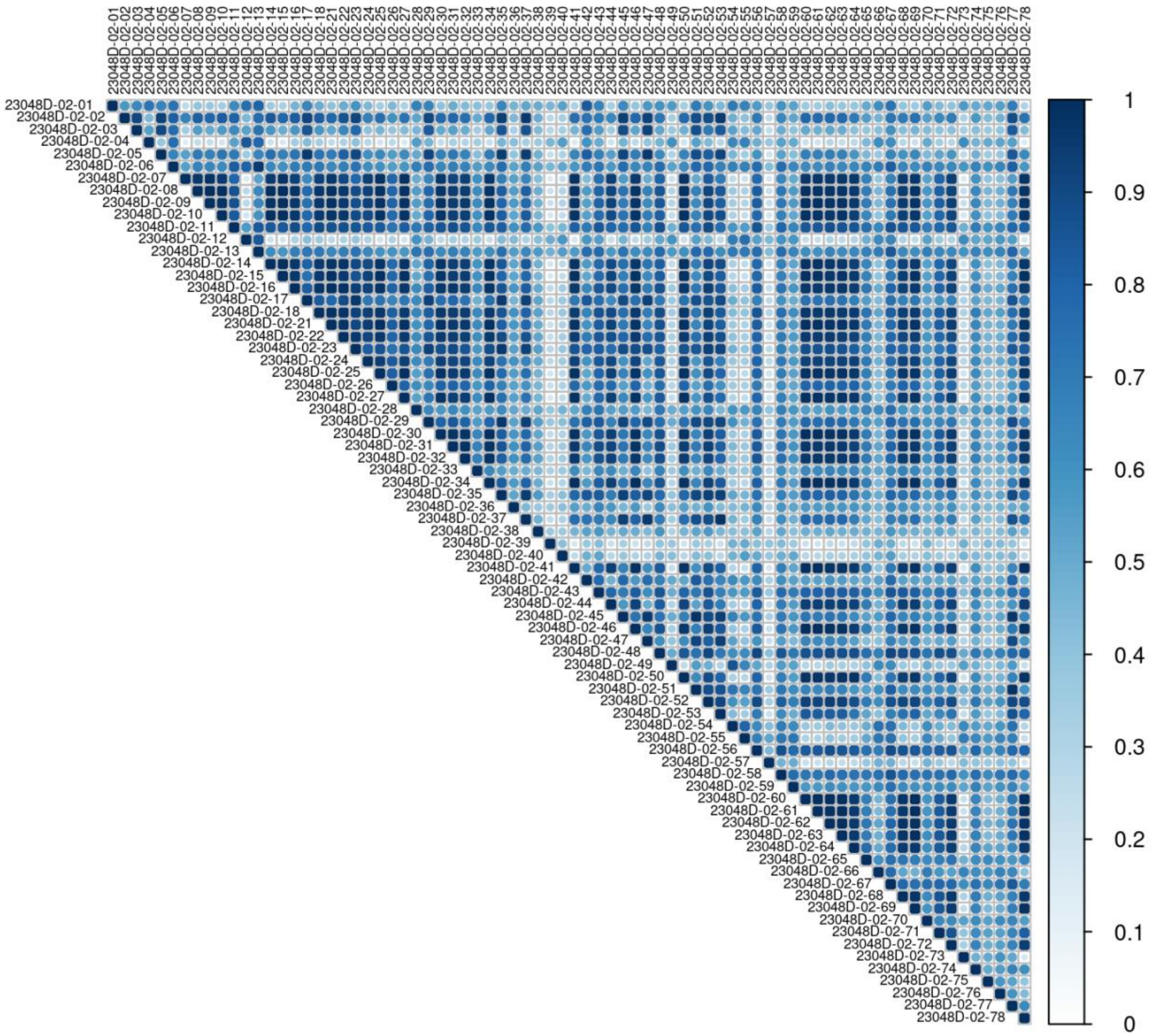


**Supplementary Figure S4**: Correlation matrix (Pearson's ρ) for pairwise comparisons of sample normalized read coverage in 1kb windows.

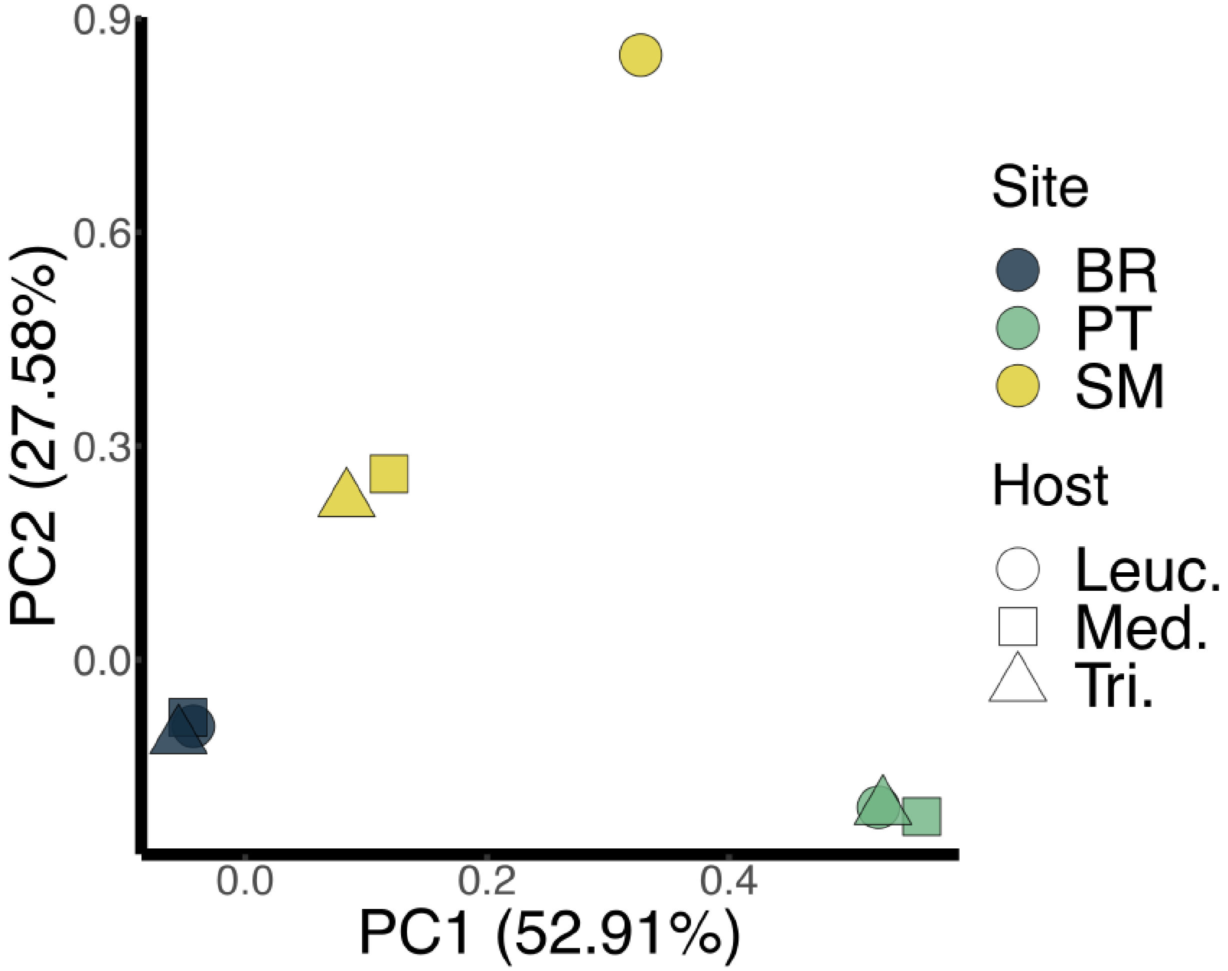


**Supplementary Figure S5:** Principal component analysis of singular-value decomposition of the scaled covariance matrix of allele frequencies **Ω**. Each point represents a host-associated group of nematodes.

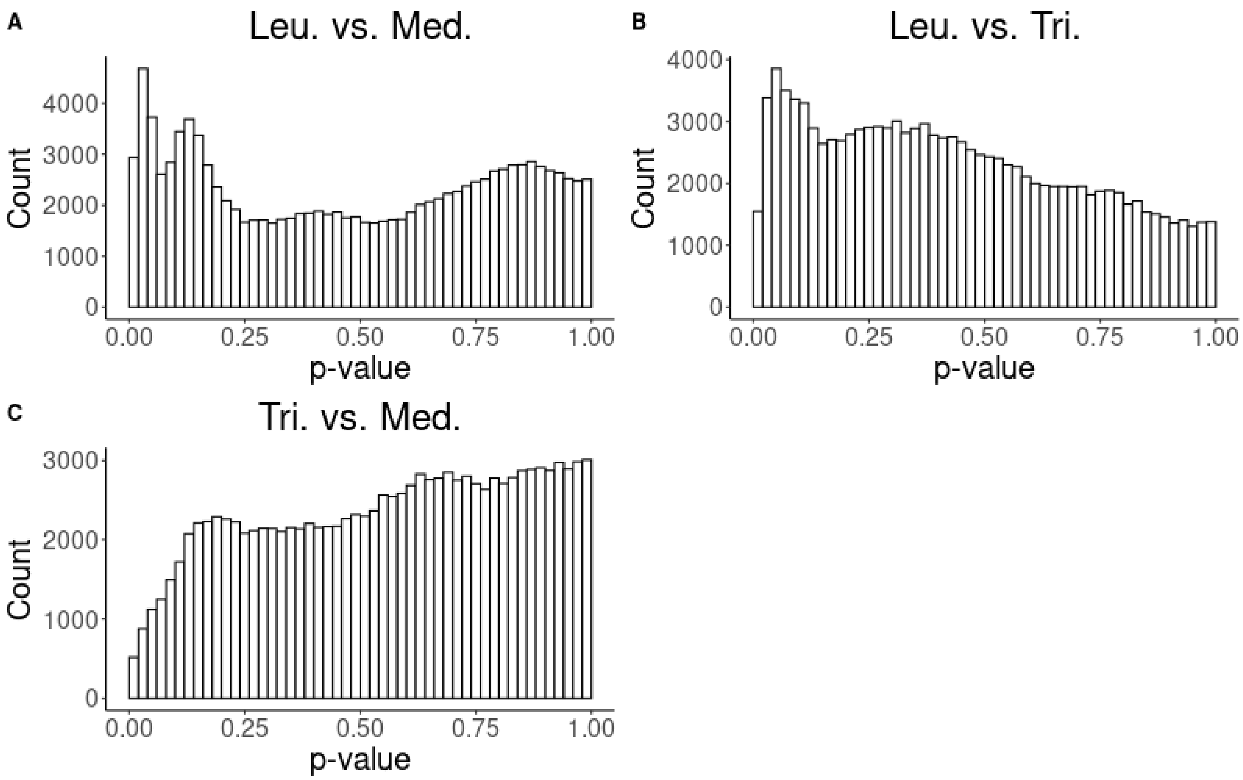


**Supplementary Figure S6**: Histogram of *p*-values for SNP-specific $C_2$ estimates for each pairwise comparison of host-associated root-knot nematode groups: (A) *L. vulgare*-associated nematodes vs. *M. lupulina*-associated nematodes (Leu. vs. Med), (B) *L. vulgare*-associated nematodes vs. *T. repens*-associated nematodes (Leu. vs. Tri), (C) *T. repens*-associated nematodes vs. *M. lupulina*-associated nematodes (Tri. vs. Med.)

**Supplementary Table S1**: Parameter estimates and parametric 90% bootstrap confidence intervals from best-fitting demographic model inferred with fastsimcoal

| **Parameter** | **Estimate** | **90% parametric bootstrap confidence interval** |
|---|---|---|
| *Effective population sizes* | | |
| Bolley Road (BR) | 28734 | 27464.83 - 30599.28 |
| Smallville Fire Road (SM) | 27038 | 25715.65 - 30987.93 |
| Pott's Mountain Trail (PT) | 15433 | 14039.95 - 17022.63 |
| *Population split times (generations)* | | |
| PT - SM | 7861 | 6911.95 - 8689.78 |
| BR - PT | 8998 | 7635.88 - 9597.78 |
| *Contemporary migration rates* | | |
| BR to SM | $2.3822e^{-6}$ | $5.4643e^{-8}$ - $3.3862e^{-6}$ |
| SM to BR | $5.4429e^{-5}$ | $4.9339e^{-5}$ - $5.7403e^{-5}$ |
| BR to PT | $1.4973e^{-4}$ | $1.3368e^{-4}$ - $1.6365e^{-4}$ |
| PT to BR | $3.9428e^{-5}$ | $3.5625e^{-5}$ - $4.363e^{-5}$ |
| SM to PT | $1.9315e^{-6}$ | $6.7123e^{-8}$ - $3.9697e^{-6}$ |
| PT to SM | $4.5841e^{-7}$ | $4.3914e^{-8}$ - $1.095e^{-6}$ |
| *Ancient migration rates* | | |
| BR to PT | $2.6594e^{-5}$ | $1.0924e^{-7}$ - $7.9387e^{-5}$ |
| PT to BR | $6.2444e^{-8}$ | $4.7252e^{-8}$ - $2.6603e^{-4}$ |
| *Population growth rates (negative values indicate positive growth)* | | |
| BR | $-1.0389e^{-4}$ | $-1.1165e^{-4}$ - $-1.0048e^{-4}$ |
| SM | $-5.9032e^{-4}$ | $-6.2222e^{-4}$ - $-5.2068e^{-4}$ |
| PT | $-4.4075e^{-4}$ | $-4.7169e^{-4}$ - $-4.0526e^{-4}$ |

**Supplementary Table S2**: Significantly enriched GO-terms for globally host-associated SNPs in regions with significantly negative Tajima's *D* values

| GO ID | Term | Annotated | Significant | Expected | P-value |
|---|---|---|---|---|---|
| GO:0016758 | hexosyltransferase activity | 46 | 3 | 0.16 | 0.00045 |
| GO:0016757 | glycosyltransferase activity | 51 | 3 | 0.18 | 0.00061 |
| GO:0000026 | Alpha-1,2-mannosyltransferase activity | 1 | 1 | 0 | 0.00354 |
| GO:0004377 | GDP-Man:Man3GlcNAc2-PP-Dol alpha-1,2-mannosyltransferase activity | 1 | 1 | 0 | 0.00354 |
| GO:0015020 | glucuronosyltransferase activity | 28 | 2 | 0.1 | 0.00406 |

**Supplementary Table S3**: Significantly enriched GO-terms for globally host-associated SNPs in regions with significantly positive Tajima's *D* values

| GO ID | Term | Annotated | Significant | Expected | P-value |
|---|---|---|---|---|---|
| GO:0016407 | acetyltransferase activity | 10 | 2 | 0.07 | 0.0019 |
| GO:0004592 | pantoate-beta-alanine ligase activity | 1 | 1 | 0.01 | 0.0067 |
| GO:0004823 | leucine-tRNA ligase activity | 1 | 1 | 0.01 | 0.0067 |
| GO:0004703 | G protein-coupled receptor kinase activity | 2 | 1 | 0.1 | 0.0134 |
| GO:0016747 | acyltransferase activity, transferring groups other than amino-acyl groups | 27 | 2 | 0.18 | 0.0136 |
| GO:0016746 | acyltransferase activity | 35 | 2 | 0.24 | 0.0223 |
| GO:0016874 | ligase activity | 36 | 2 | 0.24 | 0.0236 |

**Supplementary Table S4**: Significantly enriched GO-terms for locally host-associated SNPs in regions with significantly positive Tajima's *D* values

| GO ID | Term | Annotated | Significant | Expected | P-value |
|---|---|---|---|---|---|
| GO:0015294 | solute:cation symporter activity | 4 | 1 | 0 | 0.0029 |
| GO:0015296 | anion:cation symporter activity | 4 | 1 | 0 | 0.0029 |
| GO:0015377 | cation:chloride symporter activity | 4 | 1 | 0 | 0.0029 |

**Supplementary Table S5:** (Supplementary_table_S5.xlsx) List of known effectors and cell wall modifying enzymes. If homolog species is NA, then the gene was characterized in *M. hapla*.

**Supplementary Table S6**: Sequencing and mapping details of the 78 samples involved in this study. Coverage breadth = fraction of the reference genome with at least 1x coverage, Average coverage depth = mean number of reads per genomic position, and Eveness score = a metric, on a scale of 0-1, of the uniformity of read distribution across the reference genome.

| Sequence ID | Site | Host | Num. of reads | Mapped Reads (%) | Coverage breadth (%) | Average coverage depth | Evenness score |
|---|---|---|---|---|---|---|---|
| 23048D-02-01 | BR | L | 13,573,031 | 97.67 | 96.57 | 36.07 | 0.75 |
| 23048D-02-02 | BR | L | 16,255,426 | 97.4 | 96.87 | 43.04 | 0.77 |
| 23048D-02-03 | BR | L | 14,215,346 | 97.2 | 96.91 | 37.71 | 0.77 |
| 23048D-02-04 | BR | L | 11,689,783 | 90.31 | 96.53 | 28.83 | 0.70 |
| 23048D-02-05 | BR | L | 11,259,893 | 96.75 | 96.59 | 29.60 | 0.75 |
| 23048D-02-06 | BR | M | 13,541,100 | 97.42 | 96.64 | 35.87 | 0.73 |
| 23048D-02-07 | BR | M | 10,561,588 | 95.04 | 90.39 | 25.35 | 0.40 |
| 23048D-02-08 | BR | M | 13,339,067 | 97.82 | 96.30 | 35.13 | 0.66 |
| 23048D-02-09 | BR | M | 13,239,005 | 96.8 | 95.98 | 34.39 | 0.63 |
| 23048D-02-10 | BR | M | 13,780,035 | 96.59 | 96.73 | 35.24 | 0.62 |
| 23048D-02-11 | BR | T | 13,431,786 | 93.8 | 96.10 | 34.15 | 0.65 |
| 23048D-02-12 | BR | T | 13,232,913 | 97.55 | 96.61 | 35.02 | 0.72 |
| 23048D-02-13 | BR | T | 12,239,464 | 97.73 | 96.59 | 32.51 | 0.73 |
| 23048D-02-14 | BR | T | 11,348,340 | 97.23 | 96.35 | 29.53 | 0.66 |
| 23048D-02-15 | BR | T | 21,612,320 | 97.66 | 96.05 | 55.28 | 0.51 |
| 23048D-02-16 | PA | L | 17,191,993 | 93.48 | 98.03 | 42.50 | 0.65 |
| 23048D-02-17 | PA | L | 23,405,883 | 76.01 | 98.35 | 47.53 | 0.75 |
| 23048D-02-18 | PA | L | 21,649,156 | 96.15 | 98.10 | 55.16 | 0.70 |
| 23048D-02-19 | PA | L | 14,534,127 | 2.33 | 5.38 | 0.18 | NA |
| 23048D-02-20 | PA | M | 22,842,315 | 1.44 | 4.03 | 0.18 | NA |
| 23048D-02-21 | PA | M | 12,208,366 | 96.21 | 97.48 | 31.01 | 0.66 |
| 23048D-02-22 | PA | M | 17,565,730 | 66.59 | 97.86 | 30.76 | 0.69 |
| 23048D-02-23 | PA | M | 14,933,637 | 73.23 | 97.89 | 28.93 | 0.70 |
| 23048D-02-24 | PA | M | 13,072,866 | 87.18 | 94.26 | 30.11 | 0.53 |
| 23048D-02-25 | PA | M | 26,075,814 | 96.32 | 98.25 | 65.75 | 0.66 |
| 23048D-02-26 | PA | T | 13,526,563 | 51.95 | 77.58 | 18.37 | 0.35 |
| 23048D-02-27 | PA | T | 23,922,509 | 96.32 | 97.62 | 59.98 | 0.55 |
| 23048D-02-28 | PA | T | 23,001,509 | 84.97 | 98.21 | 51.77 | 0.68 |
| 23048D-02-29 | SM | L | 10,939,322 | 94.89 | 96.73 | 27.99 | 0.72 |
| 23048D-02-30 | SM | L | 19,914,430 | 97.52 | 96.73 | 52.28 | 0.67 |

| | | | | | | | |
|---|---|---|---|---|---|---|---|
| 23048D-02-31 | SM | L | 13,137,492 | 95.65 | 96.59 | 33.76 | 0.66 |
| 23048D-02-32 | SM | L | 11,301,679 | 98.05 | 95.84 | 29.82 | 0.62 |
| 23048D-02-33 | SM | L | 13,208,316 | 94.65 | 93.18 | 33.72 | 0.48 |
| 23048D-02-34 | SM | L | 14,656,762 | 96.67 | 96.45 | 37.90 | 0.61 |
| 23048D-02-35 | SM | L | 21,259,674 | 97.71 | 97.00 | 56.58 | 0.75 |
| 23048D-02-36 | SM | M | 21,190,514 | 97.18 | 91.85 | 55.65 | 0.48 |
| 23048D-02-37 | SM | M | 18,572,763 | 87.8 | 96.99 | 44.05 | 0.73 |
| 23048D-02-38 | SM | M | 12,870,121 | 97.47 | 93.31 | 33.86 | 0.47 |
| 23048D-02-39 | SM | M | 13,441,297 | 28.68 | 87.63 | 10.45 | 0.56 |
| 23048D-02-40 | SM | T | 13,468,101 | 97.79 | 95.26 | 35.73 | 0.59 |
| 23048D-02-41 | SM | T | 16,300,154 | 93.29 | 96.19 | 40.18 | 0.57 |
| 23048D-02-42 | SM | T | 15,139,737 | 95.89 | 96.42 | 39.42 | 0.75 |
| 23048D-02-43 | SM | T | 28,081,227 | 97.28 | 97.03 | 73.76 | 0.69 |
| 23048D-02-44 | SM | T | 15,050,966 | 92.85 | 90.69 | 37.24 | 0.42 |
| 23048D-02-45 | PT | L | 16,997,851 | 97.42 | 96.45 | 44.94 | 0.76 |
| 23048D-02-46 | PT | L | 32,450,450 | 97.76 | 96.62 | 84.72 | 0.65 |
| 23048D-02-47 | PT | L | 12,081,494 | 97.2 | 96.24 | 31.89 | 0.74 |
| 23048D-02-48 | PT | L | 12,438,501 | 97.54 | 95.66 | 32.79 | 0.68 |
| 23048D-02-49 | PT | L | 20,914,534 | 91.73 | 96.36 | 51.47 | 0.67 |
| 23048D-02-50 | PT | L | 14,593,545 | 90.61 | 96.96 | 34.86 | 0.67 |
| 23048D-02-51 | PT | L | 27,453,751 | 97.22 | 96.63 | 72.65 | 0.78 |
| 23048D-02-52 | PT | L | 16,703,216 | 97.12 | 96.41 | 43.89 | 0.75 |
| 23048D-02-53 | PT | L | 22,543,611 | 97.43 | 96.59 | 59.61 | 0.76 |
| 23048D-02-54 | PT | L | 15,018,005 | 60.57 | 96.15 | 24.75 | 0.72 |
| 23048D-02-55 | PT | L | 12,519,566 | 97.6 | 95.85 | 33.26 | 0.71 |
| 23048D-02-56 | PT | M | 13,750,325 | 97.3 | 95.76 | 36.24 | 0.69 |
| 23048D-02-57 | PT | M | 14,139,503 | 91.02 | 92.83 | 34.92 | 0.54 |
| 23048D-02-58 | PT | M | 22,285,837 | 97.67 | 96.14 | 58.36 | 0.68 |
| 23048D-02-59 | PT | M | 20,262,227 | 97.25 | 96.12 | 53.09 | 0.65 |
| 23048D-02-60 | PT | M | 25,190,857 | 97.22 | 96.36 | 65.76 | 0.68 |
| 23048D-02-61 | PT | M | 11,840,013 | 97.25 | 95.40 | 30.83 | 0.64 |
| 23048D-02-62 | PT | M | 11,447,981 | 97.16 | 95.73 | 29.95 | 0.68 |

| | | | | | | | |
|---|---|---|---|---|---|---|---|
| 23048D-02-63 | PT | M | 19,720,949 | 97.37 | 96.39 | 51.50 | 0.70 |
| 23048D-02-64 | PT | M | 8,359,247 | 95.87 | 95.56 | 21.17 | 0.64 |
| 23048D-02-65 | PT | M | 23,940,510 | 95.96 | 97.48 | 60.79 | 0.60 |
| 23048D-02-66 | PT | M | 13,514,333 | 96.31 | 95.43 | 34.99 | 0.63 |
| 23048D-02-67 | PT | T | 12,781,591 | 97.54 | 96.07 | 33.79 | 0.70 |
| 23048D-02-68 | PT | T | 14,184,397 | 96.17 | 95.81 | 36.55 | 0.63 |
| 23048D-02-69 | PT | T | 20,382,911 | 93.98 | 96.22 | 51.34 | 0.65 |
| 23048D-02-70 | PT | T | 21,060,601 | 97.72 | 96.03 | 55.63 | 0.62 |
| 23048D-02-71 | PT | T | 11,784,821 | 95.21 | 93.71 | 30.29 | 0.55 |
| 23048D-02-72 | PT | T | 12,638,785 | 94.09 | 94.65 | 31.85 | 0.57 |
| 23048D-02-73 | PT | T | 22,066,810 | 98.23 | 95.85 | 58.79 | 0.61 |
| 23048D-02-74 | PT | T | 11,691,424 | 97.77 | 95.04 | 30.88 | 0.62 |
| 23048D-02-75 | PT | T | 21,144,737 | 96.06 | 96.88 | 53.93 | 0.62 |
| 23048D-02-76 | PT | T | 12,183,545 | 95.47 | 94.92 | 31.49 | 0.63 |
| 23048D-02-77 | PT | T | 9,064,075 | 97.03 | 96.02 | 23.84 | 0.76 |
| 23048D-02-78 | PT | T | 15,094,971 | 95.86 | 95.46 | 37.98 | 0.55 |

**Supplementary Table S7**: Whole genome amplification reaction final DNA concentrations for each sample

| Sequence ID | Site | Host | REPLI-g amplified product (ng/µl) |
|---|---|---|---|
| 23048D-02-01 | BR | L | 298.00 |
| 23048D-02-02 | BR | L | 465.45 |
| 23048D-02-03 | BR | L | 289.98 |
| 23048D-02-04 | BR | L | 220.59 |
| 23048D-02-05 | BR | L | 355.82 |
| 23048D-02-06 | BR | M | 512.61 |
| 23048D-02-07 | BR | M | 758.56 |
| 23048D-02-08 | BR | M | 197.07 |
| 23048D-02-09 | BR | M | 153.96 |
| 23048D-02-10 | BR | M | 250.97 |
| 23048D-02-11 | BR | T | 517.51 |
| 23048D-02-12 | BR | T | 211.77 |
| 23048D-02-13 | BR | T | 718.39 |
| 23048D-02-14 | BR | T | 654.69 |
| 23048D-02-15 | BR | T | 243.13 |
| 23048D-02-16 | PA | L | 204.91 |
| 23048D-02-17 | PA | L | 187.27 |
| 23048D-02-18 | PA | L | 90.26 |
| 23048D-02-19 | PA | L | 217.65 |
| 23048D-02-20 | PA | M | 409.71 |
| 23048D-02-21 | PA | M | 228.43 |
| 23048D-02-22 | PA | M | 137.30 |
| 23048D-02-23 | PA | M | 140.24 |
| 23048D-02-24 | PA | M | 219.61 |
| 23048D-02-25 | PA | M | 156.90 |
| 23048D-02-26 | PA | T | 172.58 |
| 23048D-02-27 | PA | T | 294.09 |
| 23048D-02-28 | PA | T | 217.49 |
| 23048D-02-29 | SM | L | 213.73 |
| 23048D-02-30 | SM | L | 253.91 |

| | | | |
|---|---|---|---|
| 23048D-02-31 | SM | L | 571.40 |
| 23048D-02-32 | SM | L | 272.53 |
| 23048D-02-33 | SM | L | 108.88 |
| 23048D-02-34 | SM | L | 295.07 |
| 23048D-02-35 | SM | L | 282.33 |
| 23048D-02-36 | SM | M | 236.27 |
| 23048D-02-37 | SM | M | 296.05 |
| 23048D-02-38 | SM | M | 212.75 |
| 23048D-02-39 | SM | M | 401.88 |
| 23048D-02-40 | SM | T | 267.63 |
| 23048D-02-41 | SM | T | 210.79 |
| 23048D-02-42 | SM | T | 348.96 |
| 23048D-02-43 | SM | T | 297.02 |
| 23048D-02-44 | SM | T | 267.63 |
| 23048D-02-45 | PT | L | 308.78 |
| 23048D-02-46 | PT | L | 194.13 |
| 23048D-02-47 | PT | L | 244.11 |
| 23048D-02-48 | PT | L | 269.59 |
| 23048D-02-49 | PT | L | 297.02 |
| 23048D-02-50 | PT | L | 179.44 |
| 23048D-02-51 | PT | L | 258.81 |
| 23048D-02-52 | PT | L | 219.61 |
| 23048D-02-53 | PT | L | 553.76 |
| 23048D-02-54 | PT | L | 252.93 |
| 23048D-02-55 | PT | L | 159.84 |
| 23048D-02-56 | PT | M | 347.98 |
| 23048D-02-57 | PT | M | 223.53 |
| 23048D-02-58 | PT | M | 266.65 |
| 23048D-02-59 | PT | M | 282.33 |
| 23048D-02-60 | PT | M | 669.39 |
| 23048D-02-61 | PT | M | 520.44 |
| 23048D-02-62 | PT | M | 222.55 |

| | | | |
|---|---|---|---|
| 23048D-02-63 | PT | M | 243.13 |
| 23048D-02-64 | PT | M | 276.45 |
| 23048D-02-65 | PT | M | 238.23 |
| 23048D-02-66 | PT | M | 222.55 |
| 23048D-02-67 | PT | T | 244.11 |
| 23048D-02-68 | PT | T | 991.78 |
| 23048D-02-69 | PT | T | 285.27 |
| 23048D-02-70 | PT | T | 182.38 |
| 23048D-02-71 | PT | T | 169.64 |
| 23048D-02-72 | PT | T | 176.50 |
| 23048D-02-73 | PT | T | 370.52 |
| 23048D-02-74 | PT | T | 559.64 |
| 23048D-02-75 | PT | T | 246.07 |
| 23048D-02-76 | PT | T | 205.89 |
| 23048D-02-77 | PT | T | 235.29 |
| 23048D-02-78 | PT | T | 231.37 |

2019. Picard Toolkit. Broad Institute.
